# Selective cleavage of ncRNA and antiviral activity by human RNase2/EDN in a macrophage infection model

**DOI:** 10.1101/2021.08.30.458223

**Authors:** Lu Lu, Jiarui Li, Ranlei Wei, Irene Guidi, Luca Cozzuto, Julia Ponomarenko, Guillem Prats-Ejarque, Ester Boix

## Abstract

RNase2, also named the Eosinophil derived Neurotoxin (EDN), is one of the main proteins secreted by the eosinophil secondary granules. RNase2 is also expressed in other leukocyte cells and is the member of the human ribonuclease A family most abundant in macrophages. The protein is endowed with a high ribonucleolytic activity and participates in the host antiviral activity. Although RNase2 displays a broad antiviral activity, it is mostly associated to the targeting of single stranded RNA viruses. To explore RNase2 mechanism of action in antiviral host defence we knocked out RNase2 expression in the THP1 monocyte cell line and characterized the cell response to human Respiratory Syncytial Virus (RSV). We observed that RSV infection induced the *RNase2* expression and protein secretion in THP1 macrophage-derived cells, whereas the knockout (KO) of RNase2 resulted in higher RSV burden and reduced cell viability. Next, by means of the cP-RNAseq methodology, which uniquely amplifies the RNA 2’3’cyclic-phosphate-end products released by an endonuclease cleavage, we compared the ncRNA population in native and RNase2-KO cell lines. Among the ncRNAs accumulated in WT versus KO cells, we found mostly tRNA-derived fragments and secondly miRNAs. Analysis of the differential sequence coverage of tRNAs molecules in native and KO cells identified fragments derived from only few parental tRNAs, revealing a predominant cleavage at anticodon loops and secondarily at D-loops. Inspection of cleavage region identified U/C and A, at 5’ and 3’ sides of cleavage sites respectively (namely RNase B1 and B2 base binding subsites). Likewise, only few selected miRNAs were significantly more abundant in WT versus RNase2-KO cells, with cleavage sites located at the end of stem regions with predominance for pyrimidines at B1 but following an overall less defined nucleotide specificity.

Complementarily, by screening of a tRF&tiRNA PCR array we identified an enriched population of tRNA-derived fragments associated to RNase2 expression and RSV infection. The present results confirm the contribution of the protein in macrophage response against virus infection and provide the first evidence of its cleavage selectivity against ncRNA population. A better understanding of the mechanism of action of RNase2 recognition of cellular RNA during the antiviral host defence should pave the basis for the design of novel antiviral drugs.

## Introduction

Human RNase2 is a secretory protein expressed in leukocytes with a reported antiviral activity against single stranded RNA viruses [1, 2]. RNase2 is one of the main components of the eosinophil secondary granule matrix. The protein, upon its discovery, was named the Eosinophil Derived Neurotoxin (EDN), due to its ability to induce the Gordon phenomenon when injected into Guinea pigs [3–8]. Apart from eosinophils, RNase2 is also expressed in other leukocyte cell types, such as neutrophils and monocytes, together with epithelial cells, liver and spleen [8–10]. The protein belongs to the ribonuclease A superfamily, a family of secretory RNases that participate in the host response and combine a direct action against a wide range of pathogens with diverse immunomodulation properties [2, 11].

RNase2 stands out for its high catalytic activity against single stranded RNA and its efficiency against several viral types, such as rhinoviruses, adenoviruses and retroviruses, including HIV [12–14]. Recently, presence of eosinophils and their associated RNases has been correlated to the prognosis of COVID patients [15–17]. On the contrary, no action is reported against the tested bacterial species [12,18,19]. In particular, among respiratory viruses, which activate eosinophil recruitment and degranulation, the human Respiratory Syncytial Virus (RSV), which is the principal cause of death in infants [20], is probably the most studied model for RNase2 antiviral action. Indeed, RNase2 levels have predictive value for the development of recurrent wheezing post-RSV bronchiolitis [21]. RNase2 was proposed to have a role in the host response against the single stranded RNA virus [22] and early studies observed that RNase2 can directly target the RSV virion [12]. Interestingly, the protein ribonucleolytic activity is required to remove the RSV genome, but some structural specificity for RNase2 is mandatory, as other family homologues endowed with a higher catalytic activity are devoid of antiviral activity [23].

In our previous work using a macrophage infection model, we observed that RNase2 is the most abundantly expressed RNaseA superfamily member in the monocytic THP1 cell line [24]. To broaden the knowledge of the immunomodulatory role and potential targeting of cellular RNA population by RNase2 in human macrophages, we built an RNase2-knockout THP1 monocyte cell line using CRIPSR/Cas9 (clustered regularly interspaced short palindromic repeats) gene editing tool. Transcriptome of the RNase2 knockout with the unedited THP1-derived macrophage cells revealed that the top differently expressed pathways are associated to antiviral host defence (Lu et al., in preparation). Here, we explored the protein antiviral action by characterization of both THP1 native and RNase2-KO cell lines infected with RSV. The comparative infection study indicated that the knockout of RNase2 in THP1-derived macrophages resulted in a heavier RSV titre and reduced cell survival. Next, we analysed the total non-coding RNA (ncRNA) population by amplification of 2’3’-cyclic phosphate ends using the cp-RNAseq methodology and by screening a library array of tRNA-derived fragments. Results proved that RNase2 expression in macrophage correlates to a selective ncRNA cleavage pattern.

## Results

### RSV infection activated the expression of RNase2 in macrophages

RSV virus stock was obtained at a titration of 2.8×10^6^ TCID_50_/mL, as previously described [25] and THP1 macrophages were exposed to the RSV at a selected MOI of 1:1 up to 72h post of infection (poi). Following, we examined whether RSV infection induced the *RNase2* expression in THP1-derived macrophages. The *GAPDH* gene was used as a housekeeping gene control. Fig 1A shows that unstimulated macrophage cells had a constant and stable transcriptional expression of RNase2 and it was significantly upregulated in a time-dependent manner upon RSV infection. The significant *RNase2* gene levels upregulation could be detected as early as at 4h poi, with a 7-fold increase at 72 h poi. Furthermore, to determine whether the induction of *RNase2* mRNA levels correlated with an increase in protein expression, ELISA and WB were conducted to detect intracellular and secreted RNase2 protein of THP1-derived macrophages. At indicated poi time, culture medium and whole-cell extracts of macrophages infected with RSV were collected for ELISA and WB, respectively. As indicated in Fig 1B, the secreted protein levels of RNase2 was detected in human macrophages stimulated with RSV and was enhanced in response to RSV in a time-dependent manner, while no significant change of secreted RNase2 was detected in control macrophages. However, the maximum concentration of secreted RNase2 protein in macrophage culture was detected at 48h poi, with a slight decrease at 72h poi. Likewise, a similar profile was obtained by WB (Fig 1C). Taken together, our results suggest that RSV induces both RNase2 protein expression and secretion in human THP1 induced macrophages.

**Fig 1.**
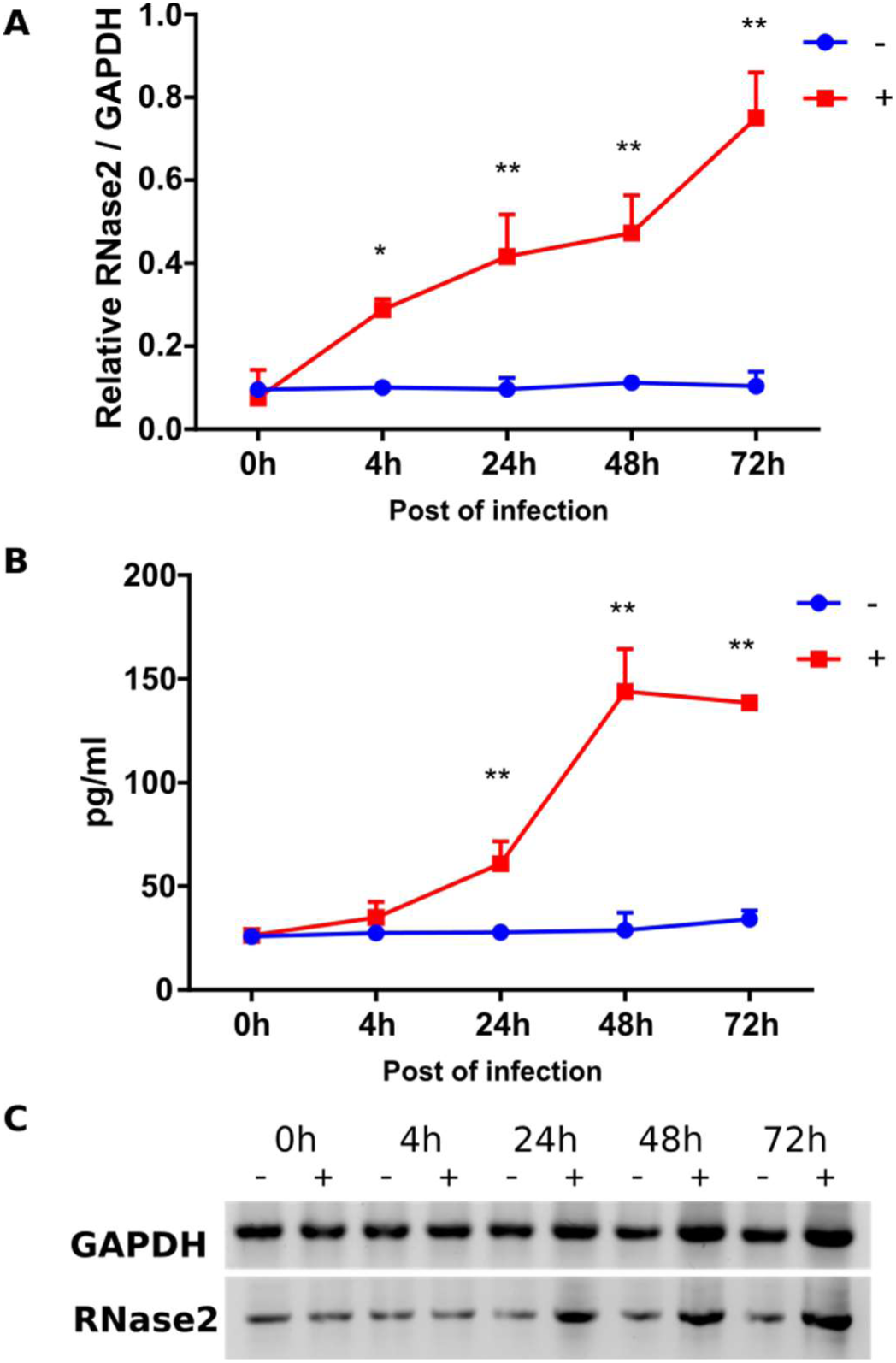
RSV infection activates the expression of RNase2 in THP1 induced macrophages. 10^6^ THP1 cells were seeded in 6-well tissue culture plate per well and induced by 50 nM of PMA treatment. After induction, macrophages were infected with RSV under MOI=1 for 2h and then cells were washed and replaced with fresh RPMI+10%FBS (0h time point post of infection). At each time point post of infection, the supernatant and cells were collected to quantify the expression of RNase2. **A)** qPCR detection of relative expression of *RNase2* gene; **B)** The concentration of the cells was controlled as 10^6^/mL, secreted RNase2 in culture supernatant was measured by ELISA and normalized with alive cell number detected by MTT assay; **C)** The intracellular RNase2 protein in macrophage was detected by WB; “+” and “-” indicate with or without RSV infection, respectively; * and ** indicate the significance of p<0.05 and p<0.01, respectively.

### RNase2 protects macrophages against RSV infection

To further characterize the protein mechanism of action, we knocked out RNase2 gene by the CRISPR/Cas9 methodology. *RNase2* gene was successfully knocked out in 2 out of 32 single cells derived THP1 cell lines, named as KO18 and KO28. Both of KO18 and KO28 express a 15aa in length peptide comparing the wild type THP1, which encodes the full length 134aa protein (Fig 2A).

**Fig 2.**
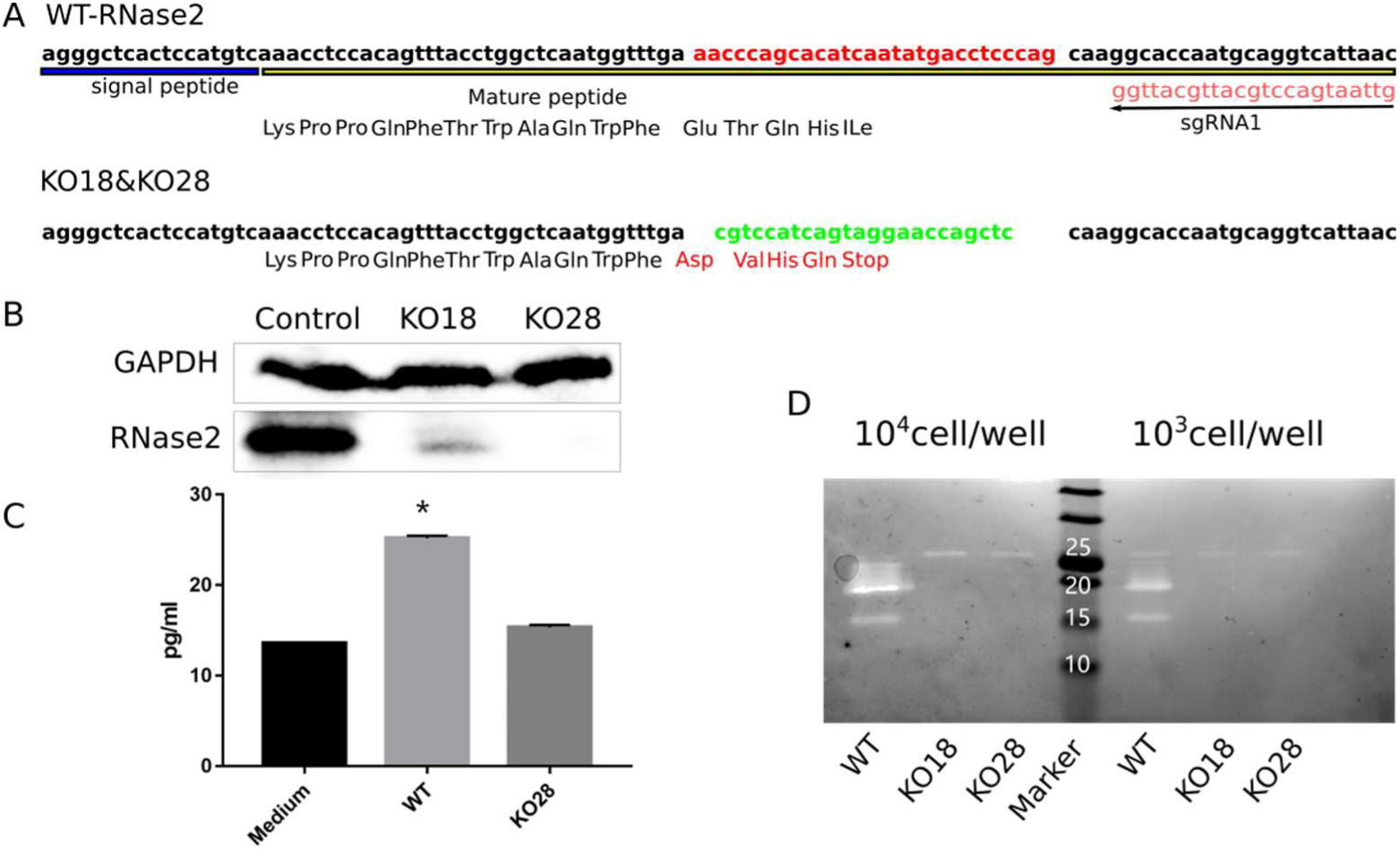
CRISPR/Cas9 mediated knock out of *RNase2* gene in THP1 derived macrophages. **A)** Scheme of the mutation of RNase2 caused by sgRNA1, the sequence was validated by Sanger sequencing; replacement is indicated: red labelled sequence in wild type was replaced by the green labelled sequence, resulting in the coding frame change and stop codon insertion; **B)** Western blot assay was applied to detect RNase2 protein; **C)** The secreted RNase2 in supernatant was measured by ELISA, the RPMI+10%FBS complete culture medium was used as a negative control, the supernatant was concentrated 50x; **D)** Ribonuclease activity staining assay was used to confirm the removal of catalytic function. Cells were collected and resuspended in water and sonicated, cell lysates were loaded in each well at the indicated quantity.

After confirmation of successful gene deletion by Sanger sequencing, we ensured that the expression of functional RNase2 has effectively been abolished. According to Western Blot (WB) assay, we can barely detect RNase2 in the KO18 cell lysate sample compared to the control sample and a total absence of signal is achieved in KO28 sample (Fig 2B). Moreover, the total absence of secreted RNase2 by THP1 cells was confirmed by ELISA assay in culture supernatant for KO28 line (Fig 2C). Last, we conducted the ribonuclease activity staining assay to evaluate the ribonucleolytic activity of the samples from cell lysates and culture supernatants. According to the activity staining electrophoresis, two main activity bands can be visualized in wild-type macrophage lysate sample, with molecular weight sizes around 15 and 20kDa, as previously reported [2]. Compared to the WT control, both activity bands were missing in the RNase2 knockout cell lines (Fig 2D). In addition, as recent studies suggested that CRISPR/Cas9 frequently induces unwanted off-target mutations, we evaluated the off-target effects on these transduced monocytes. Here, we examined the top 4 potential off-target sites for the sgRNA1 (Table S1) and did not detect off-target mutation in the T7EI assay (Fig S1). Overall, we confirmed that the RNase2 gene has been both structurally and functionally knocked out. The KO28 THP1 cell line, which achieved full RNase2-knockout, was selected for all the downstream experiments.

Next, we investigated whether RNase2 expression within macrophages contributes to the cell antiviral activity. First, THP1 cells (WT and RNase2 KO) were induced into macrophages as described above. Macrophages were then exposed to RSV at MOI=1 to investigate the kinetics of infection by monitoring both intracellular and extracellular RSV amplicon using probe RT-qPCR. Intracellularly, RSV increased at the beginning of the infection (24h) but decreased at longer infection periods (48-72h), with a slow increase (0h-4h) followed by an exponential increase (4h-24h) and reaching a maximum at 24h (Fig 3A). At 24h, RNase2 KO macrophages had significantly more intracellular RSV than WT macrophages. The extracellular RSV titre was also determined (Fig 3B). We observed that RSV increased in KO macrophage cell cultures until 48h and then stabilized at 48-72h. While in WT macrophages, RSV profile showed an increase that reached a peak at 48h and was followed by a decrease, significantly higher RSV levels were detected in KO macrophage cultures at 24h-72h. Moreover, we monitored cell death during RSV infection using MTT assay and our results confirmed that RSV infection increased cell death in either WT or KO macrophages. However, significant differences were detected, where KO macrophage cell death upon RSV infection was higher than in WT macrophage cells (Fig 3C). Altogether, we concluded that RNase2 KO macrophages burdening and cell death is significantly higher for RSV infection in comparison to WT macrophages. The present results confirm the direct involvement of the macrophage endogenous RNase2 in the cell antiviral activity.

**Fig 3.**
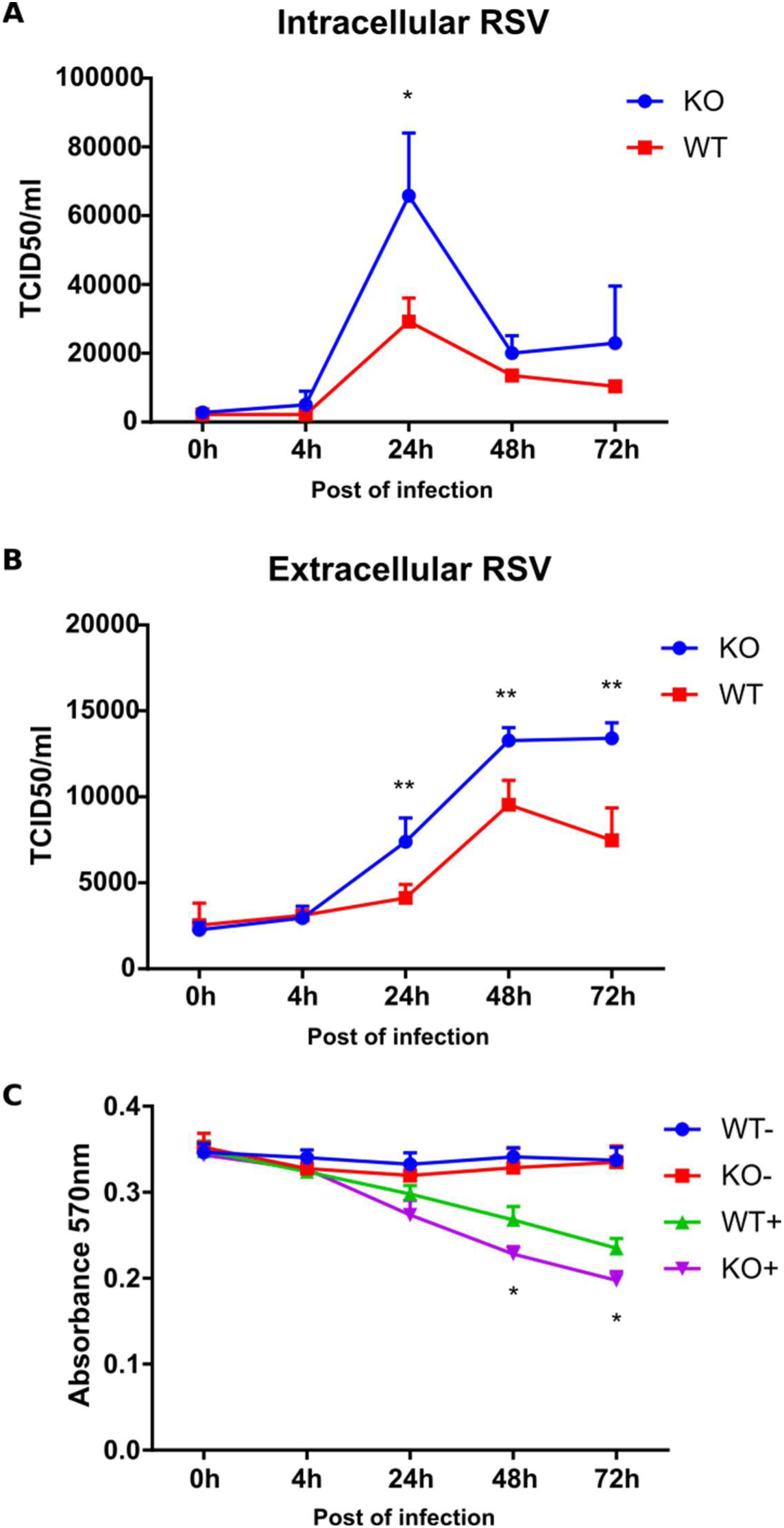
Knocking out of RNase2 resulted in macrophages suffering heavier RSV burden and more cell death. The intracellular (**A**) and extracellular (**B**) RSV was quantified by probe qPCR and calculated as the median tissue culture infectious dose (TCID50). **C**) Macrophage cell viability was monitored by registering the absorbance at 570 nm using the MTT assay; the star refers to the significance between KO+ and WT+, “+” and “-” indicate with or without RSV infection, respectively. Significance is indicated (* *p* < 0.05).

### Selective cleavage of RNase2 on ncRNA population

Following, we aimed to identify the potential changes in small RNA population associated to the expression of RNase2 by comparison of WT and KO cell lines.

Toward this end, we applied the cP-RNA-seq methodology that is able to exclusively amplify and sequence RNAs containing a 2’3′-cyclic phosphate terminus, product of an enzymatic endonuclease activity [26]. Total small RNA from WT and KO-THP1 cells was purified and the 20 to 100 nt fraction was extracted and processed as described in the methodology section. Sequence quality control indicated that for all samples more than 95% sequences achieved an average value > 30M Reads. Following RNAseq amplification, the sequence libraries were inspected by differential enrichment analysis. Principal Component Analysis (PCA) confirmed good clustering within WT and KO and appropriate discriminant power between the two groups (Fig S2).

Results revealed that RNase2 expression in THP1 cells is mostly associated to significant changes in small RNAs and in particular in tRNA fragments and miRNAs population (see Figs. S3-S5 and additional files 1-3). Overall, we observed an overabundance of only few specific tRNA-derived fragments (Fig 4, Table S2 and Fig S4) and miRNAs (Table S3 and Fig S5).

**Fig 4:**
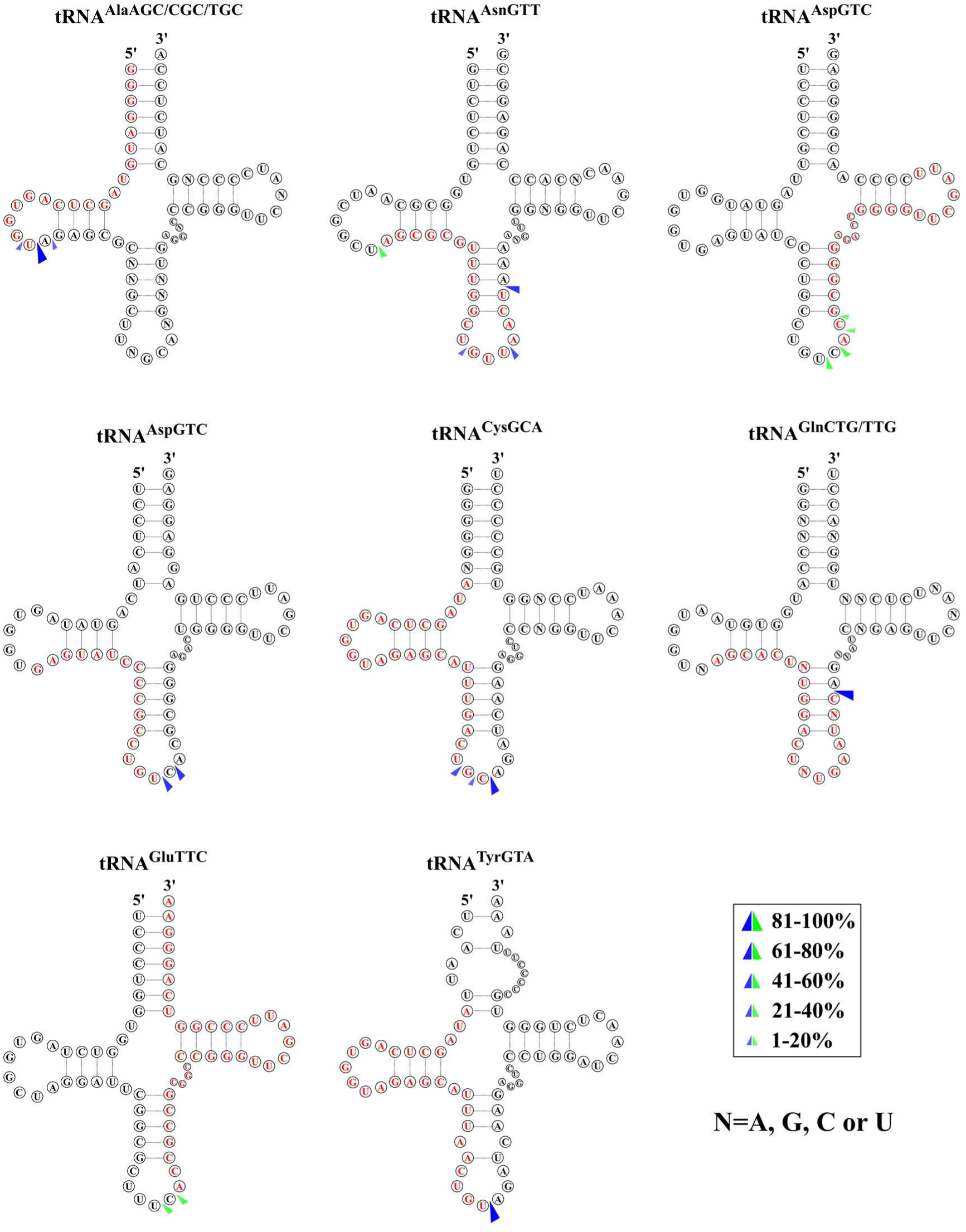
Predicted cleavage sites of the most significantly abundant tRNA-derived fragments identified by cp-RNAseq. Parental tRNAs with significant coverage differences between WT and RNase2-KO macrophage cells are depicted and the identified tRFs are marked in red. The possible cleavage sites are based on the 3’-terminal positions of the five prime fragments (blue arrows) and the 5’ terminal positions of three prime fragments (green arrows) according to the different coverage. Only sequences with a log_2_fold >0.5 and adjusted *p* value < 0.05 are included.

Analysis of differential sequence coverage between WT and KO samples (Table S2) indicated that the preferential cleavage sites for RNase2 on tRNAs were CA and UA (Fig 4).

Fig 5A illustrates the base preferences for B1 and B2 deduced from the differential sequence coverage analysis of bam files. Results highlighted a selectivity for pyrimidines at B1 and preference for purines at B2, with a U/C ≥ 1 at the 5’ side of the cleavage site, and a pronounced predilection for A at the 3’ side.

**Fig 5.**
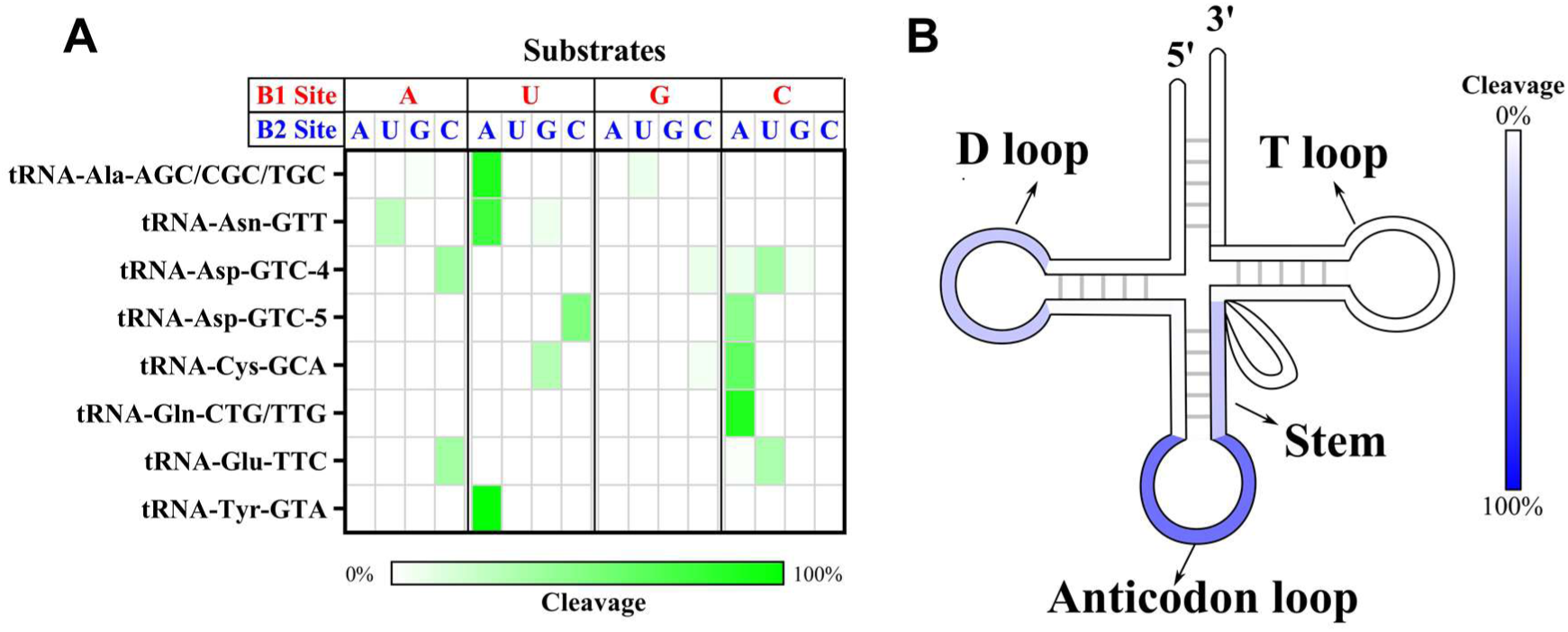
Overall cleavage preference of RNase2 on tRNAs in THP1 macrophage cells of WT vs RNase2-KO identified by cP-RNA-seq. **A**) Estimated base preference at 5’ and 3’ of cleavage site (B1 and B2 respectively) deduced from analysis of differential sequence coverage. **B**) The percentage of predicted cleavage location is depicted from low (white) to high (blue) values.

We also explored whether the cleavage preference was dependent on the RNA adopted secondary. We can see how RNase2 preferentially cuts at tRNA loops, mostly at the anticodon loop and secondarily at the D-loop, as well as stem regions near the anticodon loop (Fig 5B).

On the other hand, analysis of ncRNA products by Cp-RNAseq revealed abundance of miRNA products. Interestingly, out of the more than 2000 miRNAs in human genome there were only 14 miRNA products significantly altered, of which several derived from the same parental miRNA precursor (additional file 3). Inspection of miRNA overrepresented in WT vs RNase2-KO by comparison of sequence coverage indicated a marked cleavage preference at the end of stem regions with a less defined base preference (see Fig 6 and Table S3). Overall, we can infer for RNase2 a slight preference for U and C at B1 site, followed by G, and no defined consensus for B2.

**Fig 6.**
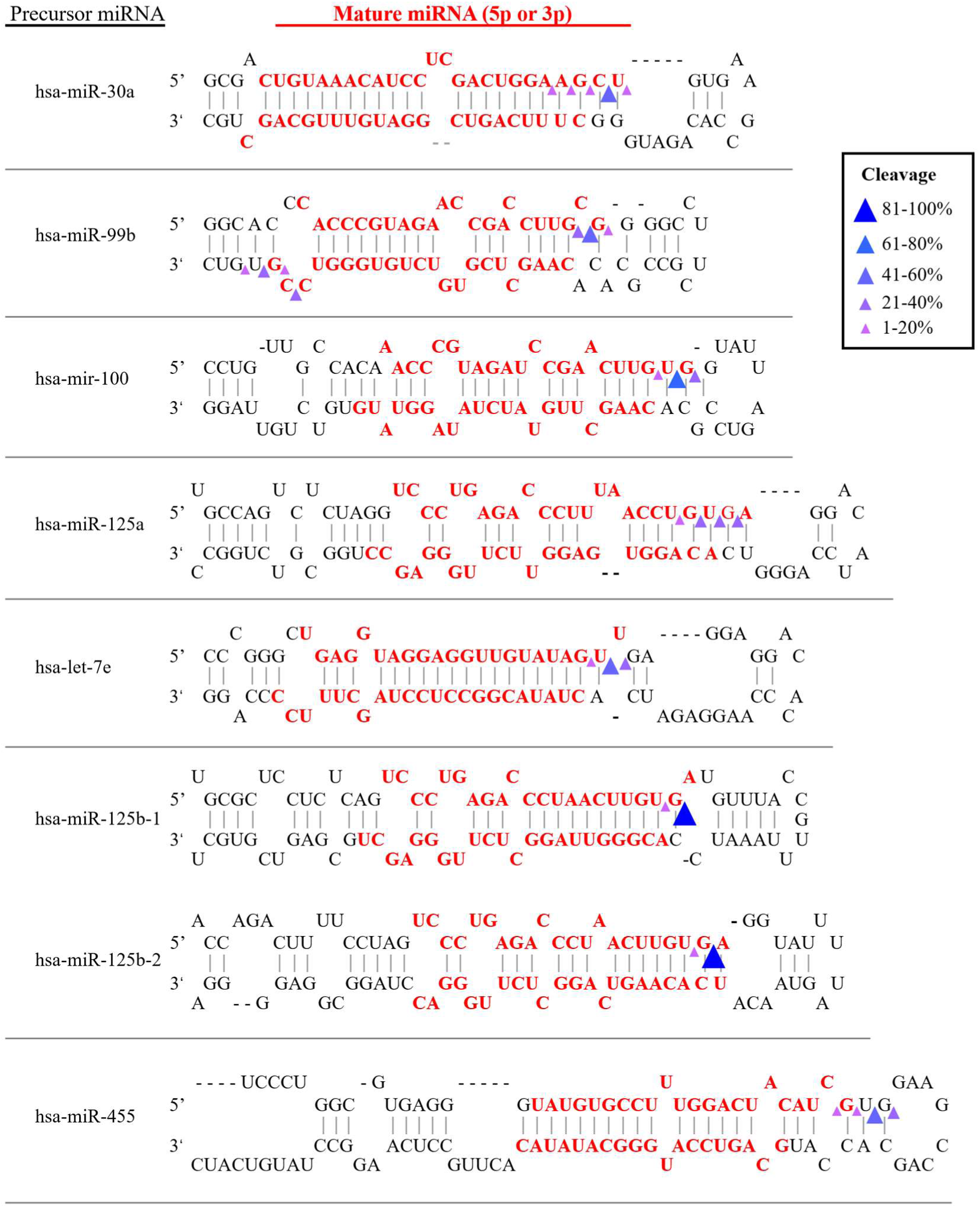
Representative miRNA overrepresented in WT versus RNase2-KO samples (log_2_fold >2 and *p* < 0.05) and predicted cleavage sites on precursor miRNA to release mature miRNAs. Cleavage sites in the precursor miRNA were predicted based on the 3’-terminal positions of each miRNA in bam files.

In parallel, we decided to explore the changes in tRNA-derived fragments population associated with RNase2 expression in THP1 cells by screening a tRNA-derived fragment library. The nrStar^TM^ Human tRF&tiRNA PCR array includes a total of 185 regulatory tRNA-derived fragments, of which 110 are taken from the tRF and tiRNA databases and 84 have been recently reported in the bibliography.

Using the nrStar^TM^ Human tRF&tiRNA library, we found that out of a total of 185 tRNA fragments, only 5 were significantly decreased in RNase2 KO macrophage in comparison to the WT control group in uninfected samples and 22 under RSV infection: 6 tiRNAs, 4 itRFs, 9 tRF-5, 4 tRF-3, 1 tRF-1 (see Table 1). Overall, the most significant changes associated to RNase2 in both infected and non-infected cell cultures (*p* < 0.01) are observed in release products from few parental tRNAs.

**Table 1.**
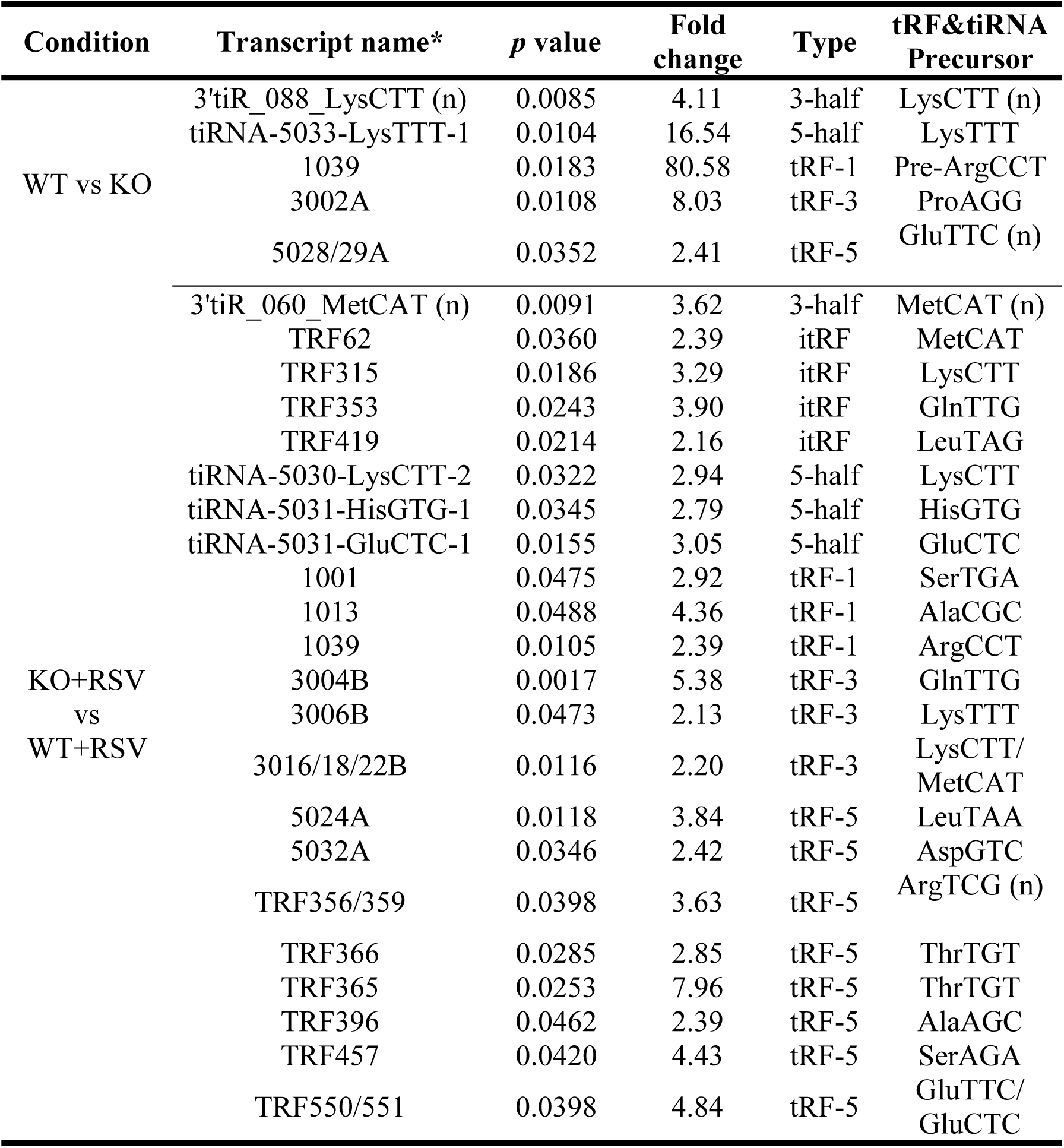
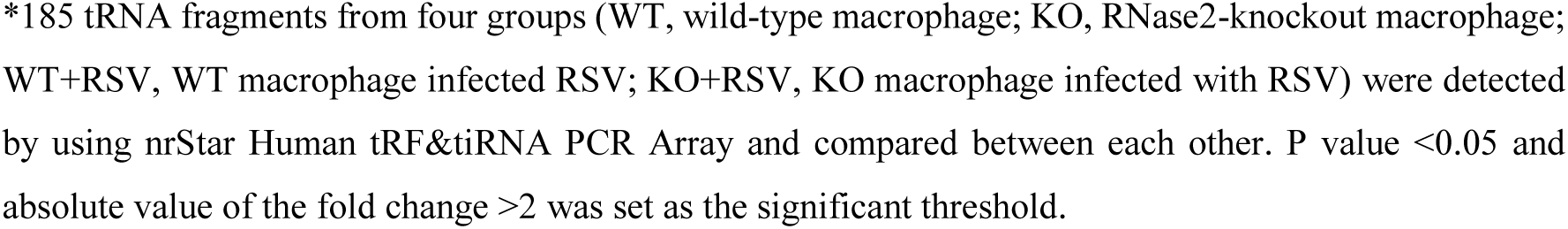
tRNA fragment population changes upon RNase2 knockout and/or RSV infection.

| Condition | Transcript name* | *p* value | Fold change | Type | tRF&tiRNA Precursor |
| --- | --- | --- | --- | --- | --- |
| WT vs KO | 3'tiR_088_LysCTT (n) | 0.0085 | 4.11 | 3-half | LysCTT (n) |
|  | tiRNA-5033-LysTTT-1 | 0.0104 | 16.54 | 5-half | LysTTT |
|  | 1039 | 0.0183 | 80.58 | tRF-1 | Pre-ArgCCT |
|  | 3002A | 0.0108 | 8.03 | tRF-3 | ProAGG |
|  | 5028/29A | 0.0352 | 2.41 | tRF-5 | GluTTC (n) |
| KO+RSV<br>vs<br>WT+RSV | 3'tiR_060_MetCAT (n) | 0.0091 | 3.62 | 3-half | MetCAT (n) |
|  | TRF62 | 0.0360 | 2.39 | itRF | MetCAT |
|  | TRF315 | 0.0186 | 3.29 | itRF | LysCTT |
|  | TRF353 | 0.0243 | 3.90 | itRF | GlnTTG |
|  | TRF419 | 0.0214 | 2.16 | itRF | LeuTAG |
|  | tiRNA-5030-LysCTT-2 | 0.0322 | 2.94 | 5-half | LysCTT |
|  | tiRNA-5031-HisGTG-1 | 0.0345 | 2.79 | 5-half | HisGTG |
|  | tiRNA-5031-GluCTC-1 | 0.0155 | 3.05 | 5-half | GluCTC |
|  | 1001 | 0.0475 | 2.92 | tRF-1 | SerTGA |
|  | 1013 | 0.0488 | 4.36 | tRF-1 | AlaCGC |
|  | 1039 | 0.0105 | 2.39 | tRF-1 | ArgCCT |
|  | 3004B | 0.0017 | 5.38 | tRF-3 | GlnTTG |
|  | 3006B | 0.0473 | 2.13 | tRF-3 | LysTTT |
|  | 3016/18/22B | 0.0116 | 2.20 | tRF-3 | LysCTT/<br>MetCAT |
|  | 5024A | 0.0118 | 3.84 | tRF-5 | LeuTAA |
|  | 5032A | 0.0346 | 2.42 | tRF-5 | AspGTC |
|  | TRF356/359 | 0.0398 | 3.63 | tRF-5 | ArgTCG (n) |
|  | TRF366 | 0.0285 | 2.85 | tRF-5 | ThrTGT |
|  | TRF365 | 0.0253 | 7.96 | tRF-5 | ThrTGT |
|  | TRF396 | 0.0462 | 2.39 | tRF-5 | AlaAGC |
|  | TRF457 | 0.0420 | 4.43 | tRF-5 | SerAGA |
|  | TRF550/551 | 0.0398 | 4.84 | tRF-5 | GluTTC/<br>GluCTC |

\*185 tRNA fragments from four groups (WT, wild-type macrophage; KO, RNase2-knockout macrophage; WT+RSV, WT macrophage infected RSV; KO+RSV, KO macrophage infected with RSV) were detected by using nrStar Human tRF&tiRNA PCR Array and compared between each other. P value <0.05 and absolute value of the fold change >2 was set as the significant threshold.

Upon inspection of sequence of the putative cleavage sites we observed most sites at or near any of the tRNA loops, with predominance of anticodon loop (∼ 50%), followed by D-loop (Table S4). Moreover, we observed that the preferred target sequences in WT samples were significantly enriched with U at the 5’ position of cleavage site, although not enough data is available for a proper statistical analysis. On the other hand, when analysing all the conditions, with and without infection, we also observed overall a significant preference for U at B1 at loop sequences. Moreover, the most significant tRNA fragments associated to RNase2 presence showed a U/C cleavage target for B1 at the anticodon loop (Table S4). On the contrary, the main base at B1 was G when the cleavage site was located at a stem region. As for the base located at the 3’ side of the cleavage target, we did not observe a clear distinct preference, with only a slight tendency for U, followed by A.

Overall, most of the parental tRNAs precursors identified by the array screening matched the identified by the Cp-RNAseq assay (> 70%), although some differences were observed in the identity of the accumulated fragments and their relative fold change. To note, few of the top listed precursors by the library screening (Lys^CTT^ and Met^CAT^) were also present in the amplified sequences by the Cp-RNAseq, but with a lower abundance (Additional file 2). On the contrary, few of the top listed fragments spotted by the later methodology were absent from the commercial library array.

Notwithstanding, we must bear in mind that the results obtained from the tRNA array screening are determined by the intrinsic library composition. The tiRNA&tRFs list is based on the previously reported tRNA fragments, which have been identified mainly by the characterization of other RNases [27]. On the contrary, by Cp-RNAseq method only the RNA by-products of an endonuclease enzymatic cleavage are amplified.

The present study is the first specific characterization of the catalytic activity of RNase2 on ncRNAs. Therefore, our data is the first report of the specific release of tRNA fragments associated to RNase2 expression.

## Discussion

Expression of human RNase2 is widely distributed in diverse body tissues such as liver and spleen together with leukocyte cells [2]. Among the blood cell types, RNase2 is particularly abundant in monocytes [11], which are key contributors to host defence against pathogens. A number of host defence-associated activities have been proposed for RNase2, mostly associated to the targeting of single stranded RNA virus infection [1]. In particular, the protein has been reported to reduce the infectivity of the human respiratory syncytial virus (RSV) in cell cultures [2,12,28].

Here, we studied RNase2 expression in THP1-derived macrophage upon RSV infection. Previous work indicated that RNase2 is the most abundant RNaseA family member expressed in this human monocytic cell line [24] (https://www.proteinatlas.org/). Viruses can manipulate cell biology to utilize macrophages as vessels for dissemination, long term persistence within tissues and virus replication [29]. In our working model, we observed how RSV enters human THP1 induced macrophages within the first 2 hours of infection. A fast proliferation of RSV takes place after 4h post of infection and titre of intracellular RSV viruses reaches the highest peak at 24h. Non-treated human THP1-derived macrophages stably expressed basal levels of RNase2 and upon 4h of RSV infection the protein transcription is significantly activated, showing a time-line correlation between RSV population and RNase2 expression. Besides, a significant increase of the secreted protein is detected by ELISA after 24h of infection, reaching a peak at 48h (Fig 1). It was previously reported that human monocyte-derived macrophages challenged with a combination of LPS and TNF-α produced RNase2 in a time-dependent manner [30]. However, we did not find any significant change of transcriptional expression of RNase2 upon *Mycobacteria aurum* infection [24]. Discrepancy of expression induction is also found for RNase2 secretion by eosinophils upon distinct bacterial infections. For example, *Clostridium difficile* and *Staphylococcus aureus* infection caused release of RNase2, while *Bifidobacteria*, *Hemophilus*, and *Prevotella* species infection did not [2]. In agreement, our previous work on THP1 derived macrophages infected by mycobacteria also discarded any induction of *RNase2* expression [24]. In contrast, using the same infection model and experimental protocol, we observe here how RSV infection significantly activate both the expression and protein secretion of *RNase2* in THP1 macrophage derived cells (Fig 1). The present study corroborates previous reports on RNase2 involvement in host response to viral infections [1, 2]. In particular, our work highlights the protein role in macrophage cells challenged by RSV infection. More importantly, we observe how the knockout of RNase2 in macrophage derived cells results in heavier RSV infection and cell death (Fig 2).

Considering previous reports on the contribution of RNase2 enzymatic activity in the protein antiviral activity [12] and the evidence that RSV infection alters the cellular RNA population, including the specific release of regulatory tRFs [31–33], we decided here to analyse the contribution of macrophage endogenous RNase2 on cellular small RNAs. Increasing data demonstrates that small noncoding RNAs (ncRNAs) play important roles in regulating antiviral innate immune responses [34–36]. In particular, ncRNAs derived from tRNAs, such as tRNA halves (tiRNAs) and tRNA-derived fragments (tRFs), have been identified and proven as functional regulatory molecules [37]. RSV infection together with other cellular stress processes can regulate the population of tiRNAs and tRFs. For example, it has been demonstrated that RSV infection and hepatitis viral infection can induce the production of tRFs and tRNA-halves, and their release has been related to RNase5 activity [32, 38]. RNase5, also called Angiogenin (Ang) due to its angiogenic properties, is one of the most well-known ribonucleases that are responsible for endonucleolytic cleavage of tRNA [39–41]. Surprisingly, the release of a specific tRF, tRF5-Glu^CTC^, which targets and suppresses the apolipoprotein E receptor 2 (APOER2), can also promote the RSV replication [32, 33]. In the present study, although the tRF5-Glu^CTC^ was not significantly changed upon RSV infection alone, we observed a significant reduction in the RNase2 knockout cell line challenged with RSV infection (Table 1). Besides, tiRNA-5034-Val^CAC^-3, the 5’half originated from Val^CAC^, is identified both in the present work (Table S4) and the previous mentioned study associated to RSV infection [32]. To note, we found that RSV infection on WT cell line induced the production of 4 tRFs in macrophages, in agreement to the previous study that indicated that RSV infection induced the release of tRFs in A549 epithelial cells [32], although most of the tRNA products differ, which may be attributed to the specific basal composition of each source cell line [42]. Moreover, tRFs production is also observed to be dependent on the specific viral infection type; for example, human metapneumovirus did not induce tRFs [32]. Interestingly, release of tRNA products is mostly associated to an antiviral defence mechanism [43]. For example, the tRF3 from tRNALys^TTT^, which stands out among the identified tRFs by our library array screening (Table 1), is reported to block retroviruses replication, such as in HIV-1, by direct binding to the virus priming binding site [44–46]. Other regulatory tRNA fragments underrepresented by RNase2-KO (such as 5’ Lys^CTT^ and Glu^CTC^ halves) (Table 1) were previously reported to be released by RSV infection [32, 33].

The present experimental data constitute the first evidence on the specific cleavage by RNase2 of cellular ncRNA. Results obtained by both cp-RNAseq and tRFs array screening indicate that RNase2 expression in macrophages is associated to the significant enrichment of selective miRNAs and tRNA-derived fragments (Figs 4 and 6; Tables S2 and S3). In addition, a higher frequency of cleavage takes place at tRNA single stranded regions, with predilection for the anticodon loop, followed by the D arm (Fig 5). Exhaustive analysis of differential sequence coverage in WT and RNase2-KO THP1 cells suggests that RNase2 preferentially targets at UA and CA sequences at tRNA loops. Recently, Bartok and co-workers reported a RNase2 selectivity for U at B1 site in synthetic RNAs [47]. Interestingly, according to Hornung and co-workers, the release of U>p ends by RNase2 would participate in the activation of TLR8 at the endolysosomal compartment and will contribute to sense the presence of pathogen RNA [48]. To note, we find a good agreement between RNase2 substrate specificity identified in the present cell assay study on tRNAs and the previously reported for synthetic single stranded oligonucleotides [49, 50] (see Table 2). However, some differences are evidenced at the miRNAs cleavage and in particular at the B2 site specificity, which does not fully match the reported on synthetic substrates. This discrepancy is also evident for the other two RNaseA family members described to release specific tRFs [51–54], i.e. RNase5/Ang and Onconase, an RNase purified from *Rana pipiens* with antitumoral properties (Table S5). Previous kinetic studies on RNaseA family cleavage preference using single stranded RNA substrates revealed a specificity for pyrimidines at the main B1 site and preference for purines at B2 [49, 50]. Among the family members, we observe distinct preferences for U vs C and A vs G at B1 and B2 sites respectively. Interestingly, RNase 2 shows a marked preference for U at B1 and A at B2 on synthetic oligonucleotides [49, 55], which mostly corresponds to the observed preference identified by Cp-RNAseq for tRNA in this study (Fig 5). Nevertheless, our analysis on tRNA cleavage sites would suggest a U/C ratio for B1 a bit lower than the estimated for some synthetic substrates (Table 2).

**Table 2.**
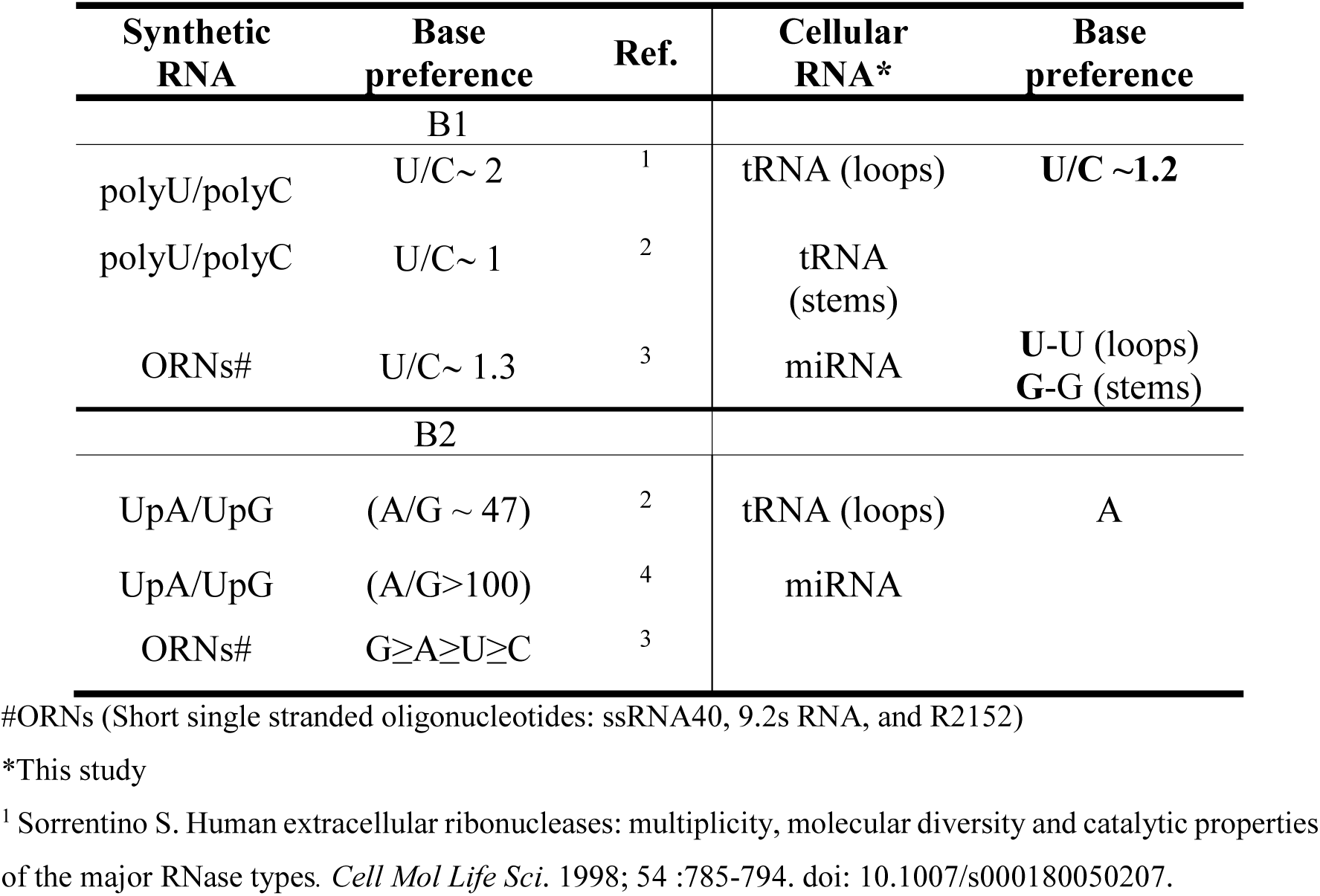
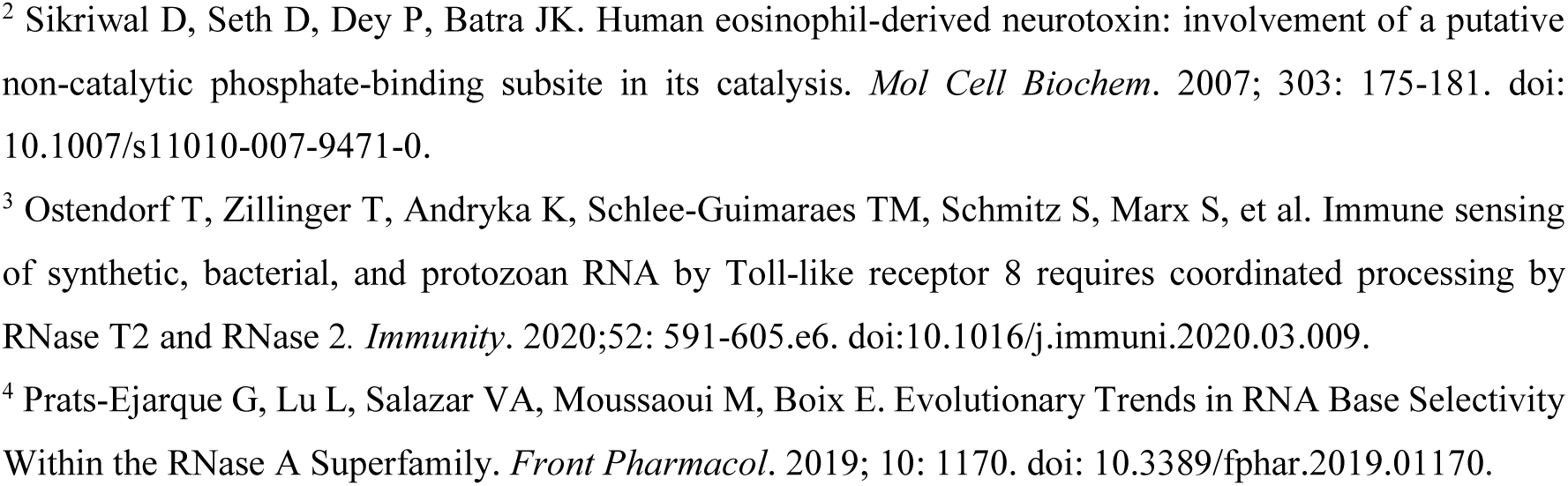
Comparison of RNase2 cleavage specificity on ncRNA in THP1-derived macrophages (this study) with synthetic RNA substrates in vitro.

| Synthetic RNA | Base preference | Ref. | Cellular RNA* | Base preference |
| --- | --- | --- | --- | --- |
| B1 |  |  |  |  |
| polyU/polyC | U/C~ 2 | <sup>1</sup> | tRNA (loops) | U/C ~1.2 |
| polyU/polyC | U/C~ 1 | <sup>2</sup> | tRNA (stems) |  |
| ORNs# | U/C~ 1.3 | <sup>3</sup> | miRNA | U-U (loops)<br>G-G (stems) |
| B2 |  |  |  |  |
| UpA/UpG | (A/G ~ 47) | <sup>2</sup> | tRNA (loops) | A |
| UpA/UpG | (A/G>100) | <sup>4</sup> | miRNA |  |
| ORNs# | G≥A≥U≥C | <sup>3</sup> |  |  |

#ORNs (Short single stranded oligonucleotides: ssRNA40, 9.2s RNA, and R2152)
\*This study
<sup>1</sup> Sorrentino S. Human extracellular ribonucleases: multiplicity, molecular diversity and catalytic properties of the major RNase types. *Cell Mol Life Sci.* 1998; 54 :785-794. doi: 10.1007/s000180050207.
<sup>2</sup> Sikriwal D, Seth D, Dey P, Batra JK. Human eosinophil-derived neurotoxin: involvement of a putative non-catalytic phosphate-binding subsite in its catalysis. *Mol Cell Biochem.* 2007; 303: 175-181. doi: 10.1007/s11010-007-9471-0.
<sup>3</sup> Ostendorf T, Zillinger T, Andryka K, Schlee-Guimaraes TM, Schmitz S, Marx S, et al. Immune sensing of synthetic, bacterial, and protozoan RNA by Toll-like receptor 8 requires coordinated processing by RNase T2 and RNase 2. *Immunity.* 2020;52: 591-605.e6. doi:10.1016/j.immuni.2020.03.009.

The present study on cellular ncRNA also highlights the key role of the RNA 3D structuration. Overall, our data reveal a cleavage preference by RNase2 at single stranded sequences and secondarily at stem adjacent to loop regions. Besides, we also observe that location of the targeted site also influences the cleavage base preference.

Interestingly, previous kinetic and structural studies on RNaseA highlighted the unusual enhancement of the protein affinity to a dinucleotide probe by addition of a phosphate linkage insert that can adopt a contorted conformation close to the cleavable 3’5’ phosphodiester bond [56–58]. Likewise, previous work on Onconase nucleotide base selectivity also encountered significant differences *in vitro* among di and tetranucleotides 1. [59] and tRNA [53].

In addition, the cleavage of tRNAs by RNases would probably be modulated by the presence of regulatory proteins within the cell [60]. For example, Onconase selectivity for specific tRNAs was attributed to the presence *in vivo* of RNA binding proteins that might protect RNA regions from RNase activity [52, 61]. In this context, we should consider the formation of regulatory complexes, such as the RISC formation, the binding of Argonaute (AGO) subunits [62] or interactions with the RNHI. On its turn, the released tRNA products would regulate the formation of cellular complexes. For example, tRF3 interaction with AGO2 mediates the cleavage of complementary Priming Binding Sequences (PBS) in retroviruses and thereby can avoid the replication of endogenous virus elements [63, 64].

Among the ncRNA population mostly altered upon RNase2 knockout, we find, together with tRFs, miRNAs. Interestingly, the identified miRNAs subproducts come frequently from the same group of parental miRNAs. Inspection on information related to these miRNAs entities reveals a predominance of miRNAs associated to cancer and neurological disorders, although few are also related to virus replication. However, caution should be taken when extracting conclusions, as miRNA databases are strongly biased from a predominance of previous clinical studies. We must also bear in mind that our working model, THP1, is a leukaemia cell line. Accumulation of miRNAs can be toxic to the cells, due to their potential interference with the translation of essential proteins. Raines and co-workers correlated miRNA release by Ang with potential toxicity to cells [41]. In any case, in our working model we do not observe any change on the viability of both WT and KO cell lines.

On the other hand, we should be aware not to over interpret our results that can be also somehow biased by the applied methodologies. The RNAseq methodology might lead to underrepresentation of some fragments, due to their relative size, short half-life or presence of posttranscriptional modifications. Therefore, caution must be taken when conclusions are drawn from the analysis of tRNA cleavage product population. Another important source of variability comes from the presence of posttranscriptional modifications, which can influence both the RNase selectivity and the product amplification step [26, 65]. Unfortunately, the current ncRNA databases are still incomplete and lack full information on the precise post-transcriptional modifications that take place *in vivo* and might intervene in the RNases recognition target.

More importantly, the array screening technique is prone to be biased by the selection criteria used to build the tiRNA&tRF array; a library composed on previously available experimental data, i.e. products by RNases such as Dicer, Angiogenin, RNaseP or RNaseZ. This might explain some of the differences observed in the identified fragments when comparing the screening of the tRFs array and the amplified sequences by the Cp-RNAseq methodology, which only amplifies the products of an endonuclease cleavage.

On the other hand, we should also take into account the protein traffic and accessibility to the distinct subcellular compartments in the assayed experimental conditions. For example, in contrast to RNase5, mostly located at the nucleus, RNase2 is associated to the endolysosomal compartment [47,48,62]. In addition, cleavage of cellular mature tRNAs might occur during stress conditions, when leakage of the RNases to the cytosol is favoured. We should bear in mind that the cell cytosolic RNA is in normal conditions protected by the presence of the RNHI, which would lose its functional conformation under stress conditions due to oxidation of surface exposed Cys residues [66]. Accordingly, in the case of RNase5/Ang, it has been described that the selective tRNA cleavage only takes place in oxidative conditions [51,67,68]. This might also explain the much higher number of tRNA fragments obtained in the present study in the RSV infected versus non-infected cells (Tables 1 and S4).

Interestingly, under certain cell conditions, such as nutrition deficiency or oxidative stress, RNase5/Ang is reported to stimulate the formation of cytoplasmic stress granules and produce tRNA-derived stress-induced RNAs (tiRNAs) [69–71]. The released tiRNAs functionally enhance damage repair and cellular survival through suppressing the formation of the translation initiation factor complex or associating with the translational silencer [40, 63]. Ivanov and co-workers recently characterized the structural determinants that guide release of tiRNA population during stress conditions [54, 72]. Accessibility of tRNAs will also depend on their potential entrapment in Tbox riboswitch or RNA granules, which are abundant in starvation situation and have an unequal propensity to protect tRNA from cleavage. Besides, proteomic analysis revealed the presence of RNHI within stress granules [73], an inhibitor protein that can complex to RNase5/Ang and other regulatory proteins to control cell translation [62].

Another important source of variability comes from the assayed cell type. Although it is widely accepted that the levels of parental tRNAs differ significantly upon cell conditions and tissues [27] and a very unequal tissue distribution is observed for the more than 500 tRNAs listed in our genome, little is still known of their relative expression rates.

Despite the inherent limitations of the present study, our results confirm that RNase2 targets ncRNA and releases specific miRNAs and tRFs. Particular interest should be drawn to the new identified tRNA fragments associated to RNase2 and absent from the commercial library array, which represent potential new regulatory elements for future studies. A growing evidence emphasizes the key role of tRNA halves and tRFs in regulating cellular functions [43,74–76]. Deciphering the contribution of the distinct RNases to shape the cell ncRNA population should be key to analyse the cell response to adapt to distinct stress conditions, such as viral infection.

## Conclusions

This is the first report of RNase2 selective targeting of ncRNA. We have engineered a THP1 cell line defective in RNase2. Comparison of native and knockout THP1-derived macrophages in a RSV infection model confirms the RNase involvement in the cell host antiviral defence. By amplification of 2’3Cp end RNA products, we have identified the tRNA fragments and miRNAs associated to RNase2 cleavage. The analysis of RNA recognition regions reveals the RNA base composition at the 5’ and 3’ of cleavage site. To note, tRNA cleavage is mostly favoured at anticodon and D-loops at UA and CA sites. Further work is mandatory for an unambiguous pattern assignment towards the understanding of how RNase2 can shape the ncRNA population and its role to fight viral infection.

## Materials and Methods

### Plasmid Construction

For long term consideration, we used the two-plasmid system to run the CRISPR gene editing experiments instead of using all-in-one CRISPR system. Thus, we constructed a pLenti-239S coding Cas9, GFP and Puromycin resistance gene for the knockout assay by replacing the sgRNA expression cassette of LentiCRISPRv2-GFP-puro (gifted by Manuel Kaulich) short annealed oligos; for activation assay, we cloned the eGFP into lenti-dCAS-VP64-Blast (Addgene61425, gifted by Manuel Kaulich), the new plasmid was named pLenti-239G. Lastly, plenti-239R, a new lentiGuide plasmid coding sgRNA expression cassette and Cherry fluorescence gene was created by using the Cherry gene (gifted by Marcos Gil García, UAB) to replace the Cas9 of LentiCRISPRv2-GFP-puro (gifted by Manuel Kaulich). The primers used for PCR are listed in Table S6.

### sgRNA design and clone into pLentiGuide (pLenti239R) vector

N20NGG motifs in the RNase2 locus were scanned, and candidate sgRNAs that fit the rules for U6 Pol III transcription and the PAM recognition domain of *Streptococcus pyogenes* Cas9 were identified. From CRISPOR (http://crispor.tefor.net/) and CRISPR-ERA, the top 2 sgRNA were selected for knockout RNase2. Using the same procedure, potential OT sites were also predicted. The sequences are listed in Table S1. Oligonucleotides were annealed and cloned into BbsI-digested pLenti-239R. The resulting plasmids containing sgRNAs were further confirmed by Sanger sequencing.

### Cell Culture

HEK293T cell line was kindly provided by Raquel Pequerul Pavón (UAB). HEK293T cell line was maintained in DMEM (Corning Life Science) supplemented with 10% fetal bovine serum (FBS) (Gibco). The culture media was replaced every 2–3 days, and the cells were passaged using Trypsin-EDTA Solution (Gibico, 25200056).

Human THP-1 cells (NCTC #88081201) were maintained or passaged in 25 or 75 cm^2^ tissue culture flasks (BD Biosciences) using RPMI-1640 (Lonza, BE12-702F) medium with 10% heat-inactivated FBS at 37°C in humidified 5% CO_2_ conditions. The culture media was replaced every 3 days.

### Generation of Lentiviral Vectors

To make the cell reach 90% confluence for transfection, 7.5×10^6^ of HEK293T cells were seeded in T75 culture flask with 15ml DMEM + 10% FBS complete medium one day before transfection. The lentiviral plasmids were transfected into HEK293T cells using calcium phosphate protocol [77]. Briefly, 36 µg transfer plasmid, 18 µg psPAX2 packaging plasmid (Addgene#35002, gifted by Marina Rodriguez Muñoz) and 18 µg pMD2G envelope plasmid (Addgene#12259, gifted by Marina Rodriguez Muñoz) were mixed. Next, 93.75 µl of 2 M CaCl_2_ was added and the final volume was adjusted to 750 µL with H_2_O. Then, 750 µL of 2×HBS buffer was added dropwise and vortexed to mix. After 15 min at room temperature, 1.5 mL of the mixture was added dropwise to HEK293T cells and the cells were incubated at 37 °C at 5% CO_2_ for 6h; then the medium was replaced with pre-warmed fresh medium. After 24h, 48h, and 72h, the supernatant was collected and cleared by centrifugation at 4000×g for 5 min and passed through 0.45 µm filter. Then, the supernatant fraction was concentrated by PEG6000 precipitate method [78] and the concentrated virus stock was aliquoted and stored at -80 °C.

### Cell Transduction

5 ×10^5^ THP1 monocytes were infected with 20 µL concentrated lentivirus in the presence of 8 µg/mL polybrene overnight. Next day, the cells were replaced with fresh medium and cultured for 72h. Fluorescence positive monocytes were checked by fluorescence microscopy and then sorted by Cell sorter. After the transduction, cells were resuspended and fixed by 2% paraformaldehyde in PBS for 10 min prior to flow cytometer. Single cells were sorted by Cell sorter BD FACSJazz.

### Sanger Sequencing

Briefly, the genomic DNA of THP1 cells was extracted using GenJET Genomic DNA purification kit (ThermoFisher, K0721) and was further used to amplify RNase2 using NZYTaq II 2×Green master mix (Nzytech, MB358). Genomic DNA was subjected to PCR (BioRad) using primers listed in Table S1. The general reaction conditions were 95°C for 10 min followed by 30 cycles of 95°C for 30 s, annealing at 60°C for 30 s, and extension at 72°C for 30 s. The pairs of primers were designed to amplify the region that covers the two possible double breaking sites. The PCR product is 410bp. After each reaction, 200 ng of the PCR products were purified using a QIAquick PCR Purification Kit (QIAGEN), subjected to T7E I assays, and then analysed by agarose gel electrophoresis. The indel mutation was confirmed by Sanger Sequencing.

### Construction of RNase2-KO THP1 cell line

A lentiviral system was selected to deliver CRISPR components into the THP1 monocytic cell line. Cas9 and sgRNAs lentiviral particles were produced in HEK293T cells by Calcium phosphate precipitation method as previously reported [77–79]. THP1 cells were then transduced with the concentrated lentiviral particles with 10 µg/mL of polybrene, the overall transduction efficiency is about 7% (Fig S6). We designed 2 single guide RNAs (sgRNAs) (Table S1) targeting the RNase2 locus to generate double strand breaks (DSBs) (Fig S7A). T7EI assay was employed to select the more active guide RNA, achieving the knockout efficiencies of about 40% (Fig S7B and S7C). The GFP and cherry red double fluorescence positive cells were then sorted by FACS into single cells and were further allowed to grow into a single cell derived clonal cell line. Following, the genomic DNA was extracted from the single cell derived cell lines and further used as a template to amplify RNase2 using the primers covering the potential mutation sites. After Sanger sequencing validation, cell lines where *RNase2* gene knock out was successful were identified.

### T7 Endonuclease I Assay-gene editing detection

As illustrated above, 200 ng of the purified *RNase2* PCR products were denatured and re-annealed in 1× T7EI Reaction Buffer and then were incubated with or without T7E I (Alt-R® Genome Editing Detection Kit, IDT). The reaction mixtures were then separated by 2% agarose gel electrophoresis. The knockout efficiency (KO%) was determined using the following formula: KO% = 100 × (1−[1−b]/[*a*+*b*])1/2, where *a* is the integrated intensity of the undigested PCR product and *b* is the combined integrated intensity of the cleavage products [80].

### Protein detection by Western Blot and ELISA

For the western blot assays, 5 ×10^5^ cells with or without transduction and their supernatants were harvested with RIPA buffer and the protein concentration was determined with the Pierce BCA Protein Assay Kit (Thermo Fisher Scientific, 23225). Equal amounts of protein (50 µg) for each sample were loaded and separated by 15% SDS-PAGE, transferred to polyvinylidene difluoride membranes. Then the membrane was blocked with 5% non-fat milk in TBST for 1 h at room temperature, and incubated with rabbit source anti-RNase2 primary antibody (abcam, ab103428) overnight at 4°C. After washing, the membranes were treated with horseradish peroxidase (HRP)-conjugated goat anti-rabbit IgG (Sigma Aldrich, 12-348) for 1 h at room temperature (RT). Finally, the membranes were exposed to an enhanced chemiluminescent detection system (Supersignal West Pico Chemiluminescent Substrate, ThermoFisher Scientific, 32209). As a control, GAPDH was detected with chicken anti-GAPDH antibodies (Abcam, ab9483) and goat anti-chicken secondary antibody (Abcam, ab6877).

Secretory RNase2 in cell culture was detected by using human RNASE2 ELISA Kit (MyBioSource, MBS773233). Beforehand, the supernatant of the culture was concentrated 50 times using 15kDa cut-off centrifugal filter unit (Amicon, C7715). Following, the standard and the concentrated culture supernatants were loaded to wells pre-coated with anti-RNase2 antibody, then the HRP-conjugated reagent was added. After incubation and washing for the removal of unbound enzyme, the substrate was added to develop the colourful reaction. The colour depth or light was positively correlated with the concentration of RNase2. Triplicates were performed for all assays.

### Zymogram/ Ribonuclease activity staining assay

Zymograms were performed as previously described [81]. 15% polyacrylamide-SDS gels were casted with 0.3 mg/mL of poly(U) (Sigma Aldrich, P9528-25MG). Then, cells were collected by centrifugation and resuspended in 10^6^ cells/ml with 1% SDS buffer. After sonication, cell lysate with indicated number of cells was loaded using a loading buffer that does not contain 2-mercaptoethanol. Electrophoresis was run at a constant current of 100 V for 1.5 h. Following, the SDS was removed by washing with 10 mM Tris/HCl, pH 8, and 10% (v/v) isopropanol for 30 min. The gel was then incubated for 1 h in the activity buffer (100 mM Tris/HCl, pH 8) to allow ribonuclease digestion of the embedded substrate and then stained with 0.2% (w/v) toluidine blue in 10 mM Tris/HCl, pH 8, for 10 min. Positive bands appeared white against the blue background after distaining.

### RSV production

Human respiratory syncytial virus (RSV, ATCC, VR-1540) stock was ordered from ATCC. Hela cells were used to produce RSV under biosafety level II conditions [82]. Briefly, Hela cells were plated in 75 cm^2^ culture flask and incubated at 37°C degree in DMEM+10%FBS until they were approximately 50% confluent. The cells were then washed and infected with RSV stock under multiplicity of infection (MOI) of 0.1. After 3h infection, the cells were washed and replaced with fresh medium (DMEM+10%FBS) and incubated for 4 days at 37°C, 5% CO_2_. The cells and the virus suspension were collected when the cytopathology appeared, with scraping and vortexing of the cells to release more viral particles. The virus suspension was centrifuged for 10 min at 1800×g to remove the cell debris. The virus suspension without cell debris were either frozen immediately and stored at -80°C as seeding stock and concentrated before use with Ultra15 Amicon 100 kDa cut-off filters. The produced viruses were titrated using the median tissue culture infectious dose (TCID50) method in HEK293T cells [83].

### RSV infection THP1 induced macrophage

Before RSV infection, THP1 cells were induced to macrophage by 50 nM of PMA treatment for 48h as previously described [24]. Cells were washed three time with pre-warmed PBS and replaced with fresh RPMI+10%FBS medium for 24h incubation. After that, macrophages were washed and incubated with RSV, mixing at every 15 min for the first 2h. All virus treatment tests were performed using RSV at a MOI of 1 TCID50/cell.

### Real-time quantitative PCR

RSV quantification were detected by real-time quantitative PCR. After the indicated time post of infection, the extracellular RSV virus were collected by PEG6000 precipitation method and intracellular RSV virus were collected by lysing the macrophage cells with the lysis buffer from mirVanaTM miRNA Isolation Kit (Ambion, Life Technologies, AM1560). Total RNA from RSV infected macrophage cells as well as stock virus was extracted using mirVanaTM miRNA Isolation Kit according to the manufacturer’s instructions. cDNA was synthesized using iScriptTM cDNA Synthesis Kit (Bio-Rad, 170-8891). The synthesis was performed using random hexamers, starting with 1 μg of total cell RNA. The RT-qPCR was performed using ddPCR ^TM^ Supermix for Probes (Bio-Rad, 1863024). Samples with a cycle threshold value of more than 40 were recorded as negative. A standard curve was prepared using serially diluted RNA extracts from a known quantity and used to quantify RSV as TCID50/mL. In parallel with the RSV probe assays, an endogenous glyceraldehyde-3-phosphate dehydrogenase (GAPDH) control was used for relative quantification of the intracellular virus. The relative expression of GAPDH and RNase2 gene in macrophages was quantified by real-time PCR using iTaq Universal SYBR Green Supermix (Bio-Rad, 1725120). The primers and probe [84] used were listed in Table S6.

### Cell viability assay

THP1 monocytes (wild type or RNase2 knockout) were seeded at 5×10^4^ cells/well in 96 well plates and differentiated into macrophages as described [24]. After infection with RSV under MOI=1 for different times, dynamic cell viability was measured using MTT assay.

### Selective amplification and sequencing of cyclic phosphate-containing RNA (cP-RNA-seq) and data analysis

Selective amplification and sequencing of cyclic phosphate-containing RNAs was performed as previously reported [26]. Briefly, small RNAs (<200nt) were extracted using mirVanaTM miRNA Isolation Kit (Ambion, Life Technologies, AM1560) as described by the manufacturer. Following RNA extraction, 20- to -100nt RNAs were purified from 8% TBE-PAGE gel. Then, the purified RNAs were treated by calf intestinal alkaline phosphatase. After phenol-chloroform purification, the RNAs were oxidized by incubation in 10mM NaIO_4_ at 0 ℃ for 40 min in the dark, followed by ethanol precipitation. The RNAs were then treated with T4 PNK. After phenol-chloroform purification, directed ligation of adapters, cDNA generation, and PCR amplification were performed using the Truseq Small RNA Sample Prep Kit for Illumina (NewEngland Biolabs, E7335S) according to the manufacturer’s protocol. The amplified cDNAs were sequenced using Illumina hiSeq2500 system at the *Centre for Genomic Regulation*, CRG, Barcelona).

For small RNA-seq analysis, skewer (v0.22) was used to remove the 5’adaptor sequences and discard low-quality reads [85]. Reads have been size selected before being aligned to the reference genome (GRCh38) with shortStack based on bowtie1 aligner [86, 87]. The mapped reads were counted with HTSeq-count [88] using the annotation from miRBase version 22.1 ((http://www.mirbase.org/) [89]. For differential analysis, DESeq2 [90] was used on count matrices of tRNA-derived fragments and miRNAs. Quantification of differential abundance of small RNAs was estimated by tRAX (http://trna.ucsc.edu/tRAX/). The selective cleavage was identified by comparison of the differential coverage in WT and KO samples. Complementarily, small RNAs were mapped based on NCBI (https://www.ncbi.nlm.nih.gov/) and Gencode 38 (https://www.gencodegenes.org/). The Integrative Genomics Viewer (IGV) was used for visualizing the aligned bam file reads and check the nucleotide composition at the putative cleavage site, which was estimated from the analysis of the differential sequence coverage of WT and RNase2-KO samples.

### tRF&tiRNA PCR Array

Total RNA was extracted by using mirVana miRNA Isolation Kit (ambion, AM1556). Next, 2 µg of purified total RNA was used to create cDNA libraries from small RNAs for qPCR detection using rtStar First-Strand cDNA Synthesis Kit (Arraystar, AS-FS-003). This method sequentially ligates 3’-adaptor with its 5’-end to the 3’-end of the RNAs, and 5’-adaptor with its 3’-end to the 5’-end of the RNAs. After cDNA synthesis, 185 (tRNA-derived fragments) tRFs & (tRNA halves) tiRNAs were profiled and quantified by qPCR method using nrStar Human tRF&tiRNA PCR Array (Arraystar, AS-NR-002).

## Acknowledgement

We thank Yundong Peng (Max Planck Institute for Heart and Lung Research, Bad Nauheim, Germany), Manuel Kaulich (Goethe University Frankfurt, Germany) for technical supporting with CRIPSR design. We would also like to show our gratitude to Laura Tussel, Anna Genesca, Marina Rodriguez Muñoz, and David Soler from UAB for helping in lentiviral production. We heartily thank Prof. Helene Rosenberg for helpful discussions.

## Funding

This research was founded by Research work was supported by the *Ministerio de Economía y Competitividad* (SAF2015-66007P and PID2019-106123GB-I00), co-financed by FEDER funds, and by *Fundació La Marató de TV3* (2080310). LL and JL were supported by *China Scholarship Council (CSC)* predoctoral fellowships.

## Notes

### Competing Interest Statement

The authors have declared no competing interest.

