## Supplemental figures and Tables for "Selective cleavage of ncRNA and antiviral activity by human RNase2/EDN in a macrophage infection model"

[4]Bioinformatic Unit. Centre de Regulació Genòmica (CRG), Barcelona, Spain

*Both authors contributed equally to this work

#### List of additional files

**Additional file 1.** List of all small RNAs identified by differential analysis of WT vs RNase2-KO THP1-derived macrophages using Cp-RNAseq methodology.

**Additional file 2.** List of tRNA-derived fragments identified by differential analysis of WT vs RNase2-KO THP1-derived macrophages using Cp-RNAseq methodology.

**Additional file 3.** List of miRNAs identified by differential analysis of WT vs RNase2-KO THP1-derived macrophages using Cp-RNAseq methodology.

### Supplementary figures

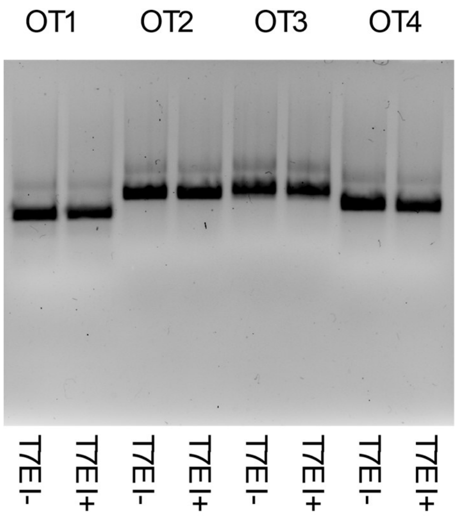


**S1 Fig. Analysis of the potential off-target effects of sgRNA1.** The sgRNA1 mediated off-target sites OT1-4 were predicted and analysed by T7EI assays.

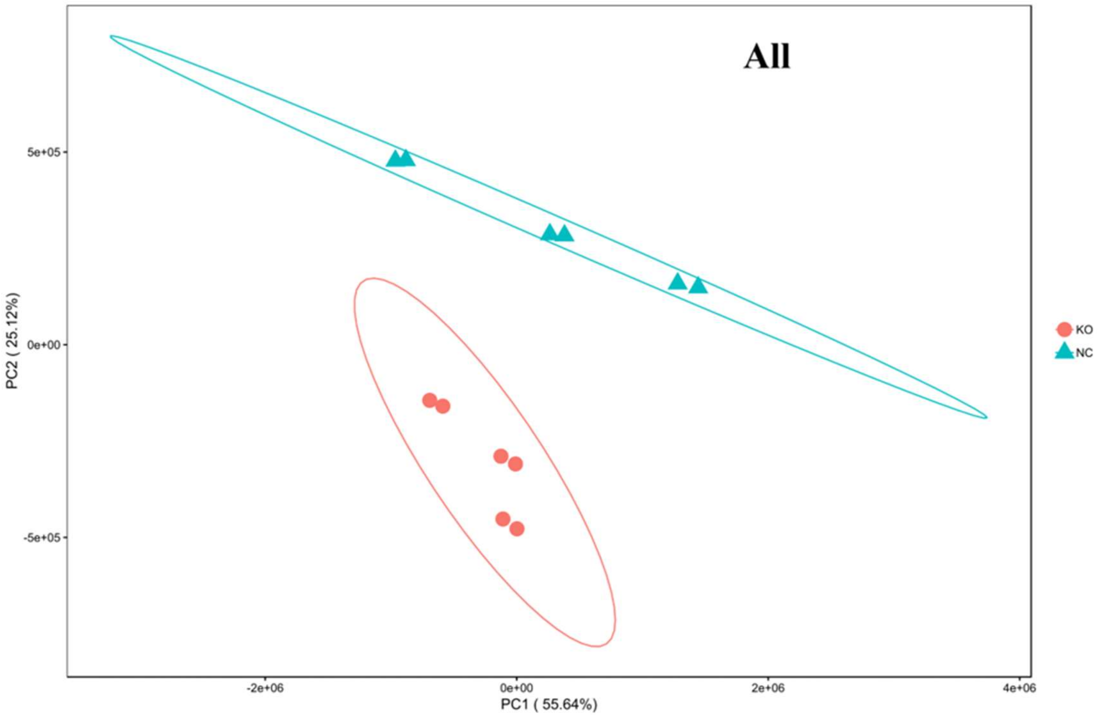


**S2 Fig. Principal analysis component (PCA) for THP1 WT and RNase2-KO for all ncRNA analysis by Cp-RNAseq.**

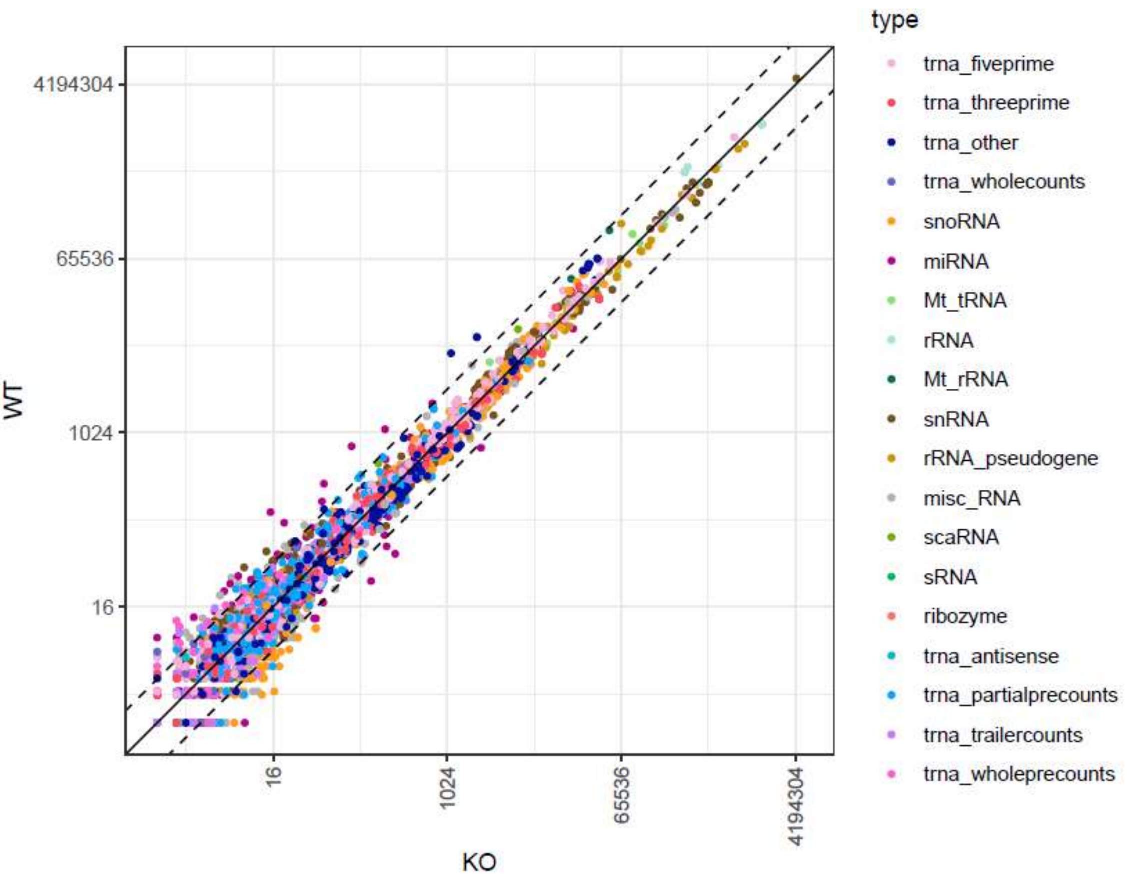


**S3 Fig. Small RNA type scatter plot for WO vs RNase2-KO samples**. For a full list of small RNAs identified by Cp-RNAseq and statistics see Additional file 1.

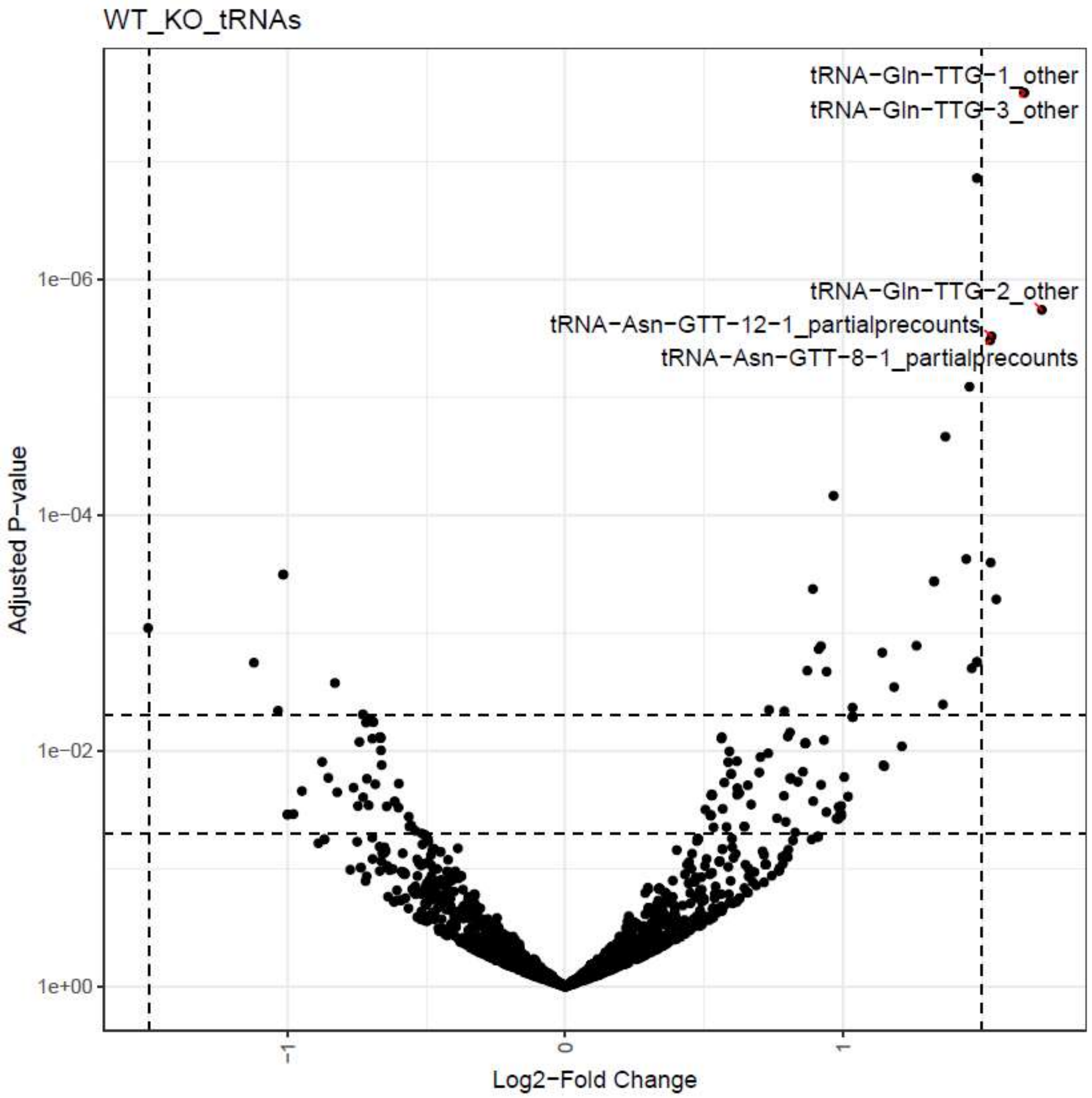


**S4 Fig. Volcano Plot for tRNAs and tRNA-derived fragments in THP1-WT vs KO cells.** The significantly down-regulated and up-regulated tRNA fragments identified by Cp-RNAseq are shown (cut-off values of Log2FoldChange and padj are indicated). The top significant tRNA fragments are labelled with their name. See Additional file 2 for a complete list of sequences and statistics.

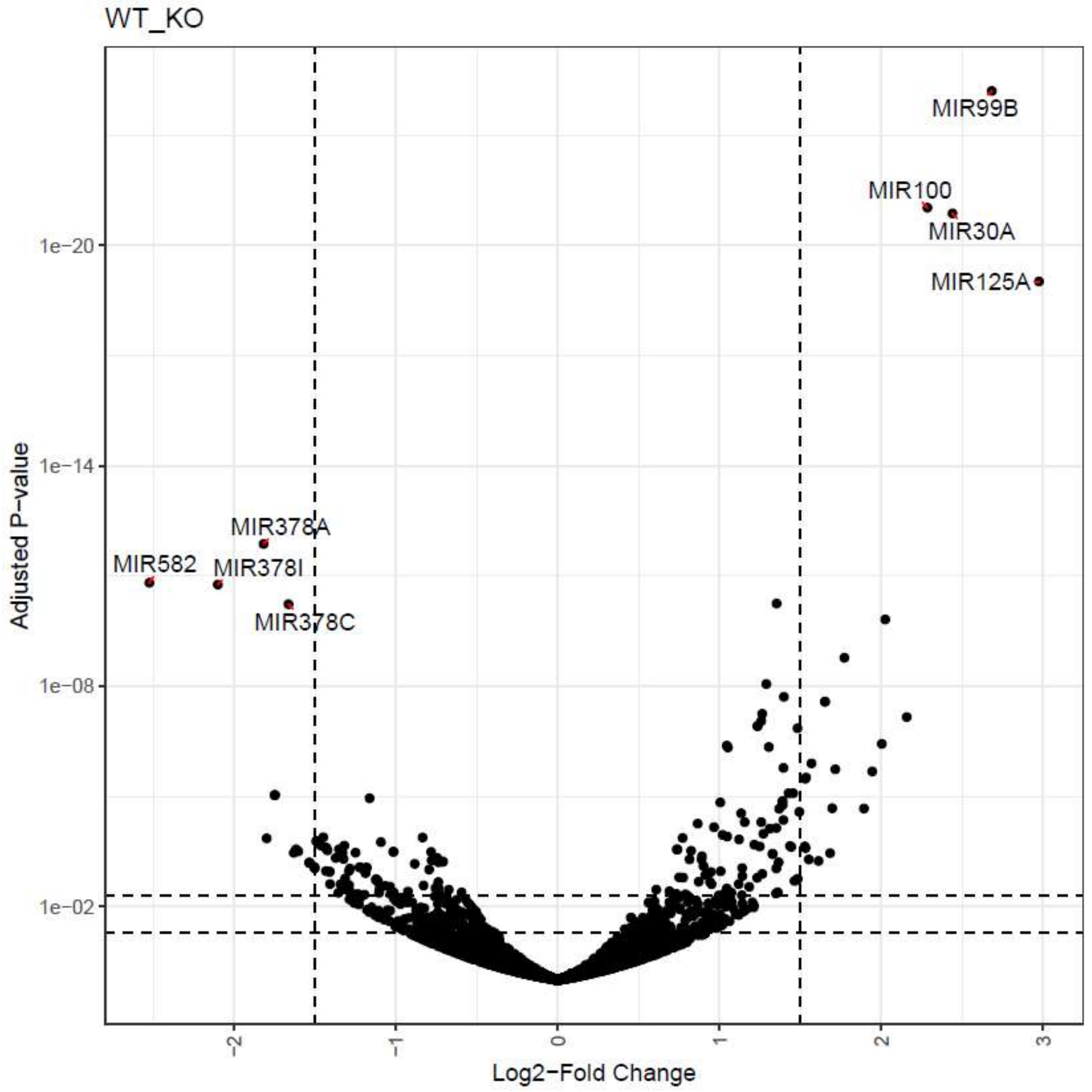


44

45 **S5 Fig**. **Volcano plot for miRNAs abundance in THP1-WT vs KO cells identified by**
46 **Cp-RNAseq methodology.** See Additional file 3 for a complete list of sequences and
47 statistics.

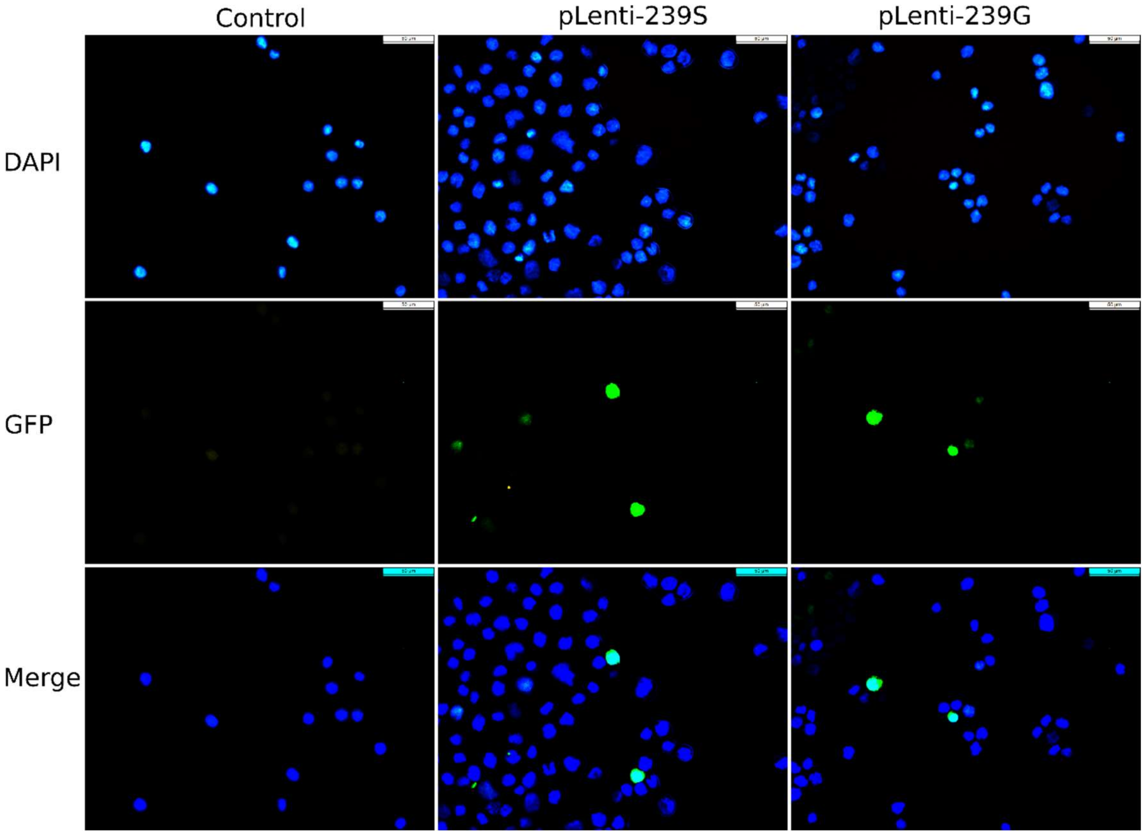


**S6 Fig. Representative image of THP1 cells tranduced by CRISPR lentiviral.** 1x$10^6$ of THP1 cells were resuspended in 2mL of RPMI+10%FBS and incubated in 37 ℃ at 5% $CO_2$ for 2h before the transduction. 20 µl of the concentrated virus and 8 µg/mL polybrene were added into THP1 culture in 6-well plate. After overnight's culture, the medium was changed with fresh medium and the cells were continually grown. After 72h post of the infection, the cell samples were fixed by 4% paraformaldehyde and dyed with DAPI and the fluorescence images were captured using Olympus BX50 fluorescence microscopy. The GFP positive cells were calculated to evaluate the transduction efficiency of lentiviral with THP1 cells from 15 randomly chosen images.

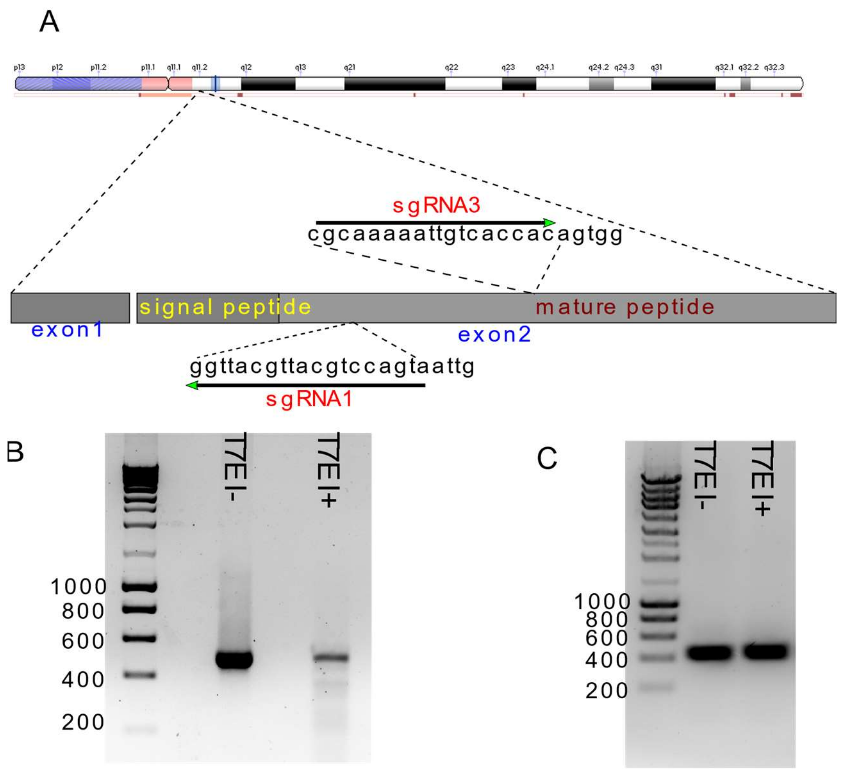


**S7 Fig. Evaluation of the knockout efficiency of sgRNA1-6 targeting the AAVS1 locus in HeLa cells. A)** Schematic of sgRNA1&3 designed to target the RNase2 locus; **(B)** and **(C)** Detection of the knockout efficiency of sgRNA1 and sgRNA3 using a T7E I assay, respectively. Amplicons of sgRNA1&3 that were treated with T7E I (T7E I+) or without T7E I (T7E I+) were separated by 2% agarose gel electrophoresis. DSB sites were recognized and digested by T7E I. Undigested and digested bands were consistent with the predicted sizes from the RNase2 locus.

### Supplementary tables

**S1 Table. Oligos of the sgRNAs and sequencing primers**

| Target name | Sequence | |
|---|---|---|
| SgRNA1 | GTTAATGACCTGCATTGCATTGG | |
| sgRNA3 | CGCAAAAATTGTCACCACAGTGG | |
| **Off-target name** | | Chromosome location |
| OT-1 | GTTAATGACCTGCATTGCATTGG | chr10:116445782-116445804:+ |
| OT-2 | CTTAATGATATGCATTGCAT AGG | chr4:95875569-95875591:- |
| OT-3 | GATAATGAGATACATTGCAT TGG | chr2:211644633-211644655:+ |
| OT-4 | TTTTATGACCTGTATAGCAT AGG | chr6:22605178-22605200:+ |
| **PCR primers** | | |
| sgRNA-x-Fw | TCTTCTGTTGGGGCTTCTGGCTG | 482 |
| sgRNA-x-Rv | GATGAGTGATGATGAGGAGTGCT | |
| sgRNA3-Fw | TCTTCTGTTGGGGCTTCTGGCTG | 482 |
| sgRNA3-Rv | GATGAGTGATGATGAGGAGTGCT | |
| OT1-Fw | TCAGGGGCTAAAATAGGGTGC | 107 |
| OT1-Rv | TATTGGATTGCCTGCTCCCC | |
| OT2-Fw | ACAAACTCTGCAGCTTTGGC | 123 |
| OT2-Rv | TCTATGTAAACAGAGTCTTTGGCA | |
| OT3-Fw | AGGGGCTTGACTCAGAGAGT | 129 |
| OT3-Rv | GGCATGCATCCATTGAGCTG | |
| OT4-Fw | TCCATGTAATCTGCCAGCCA | 115 |
| OT4-Rv | AATTGGGTGGCTGAGGTGAC | |
| **Sequencing primers** | | |
| U6-seq | GAGGGCCTATTCCCATGATT | |
| Fw-seq | GCTTCTTCTGTTGGGGCTTC | |
| Rv-seq | TCAACGACGAGACCCTCCAC | |

**S2 Table. List of parental tRNAs with significant differential coverage between WT and RNase2-KO macrophage cells.** Predicted cleavage sites of the most significantly abundant tRNA-derived fragments identified by cp-RNAseq (see Additional file 2 for a full list) is indicated. Only sequences with a $log_2$fold >0.5 and $p$ adjusted < 0.05 are included.

| Classification | $log_2$ (WT/KO) | $p$ (WT/KO) | Main predicted cleavage site | Differential coverage between WT (black) and KO (grey) |
|---|---|---|---|---|
| tRNA-Ala-AGC-4 | 0.53 | 0.0239 | GU AG | 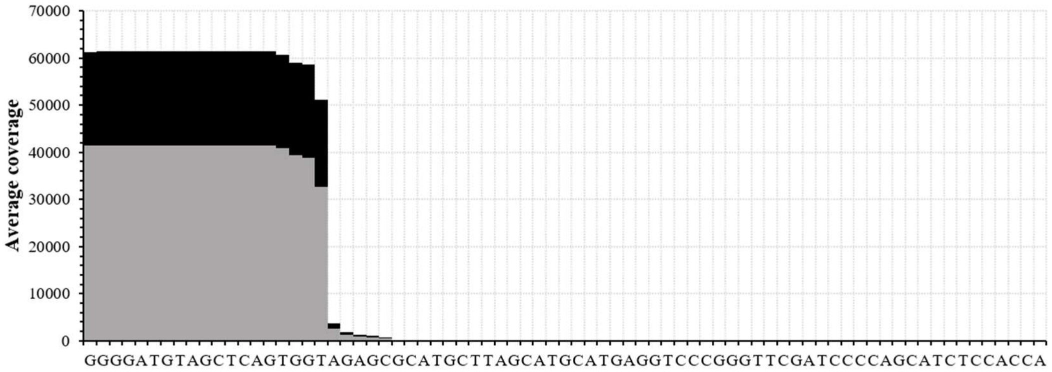 |
| tRNA-Ala-CGC-1 | 0.53 | 0.0239 | GU AG | 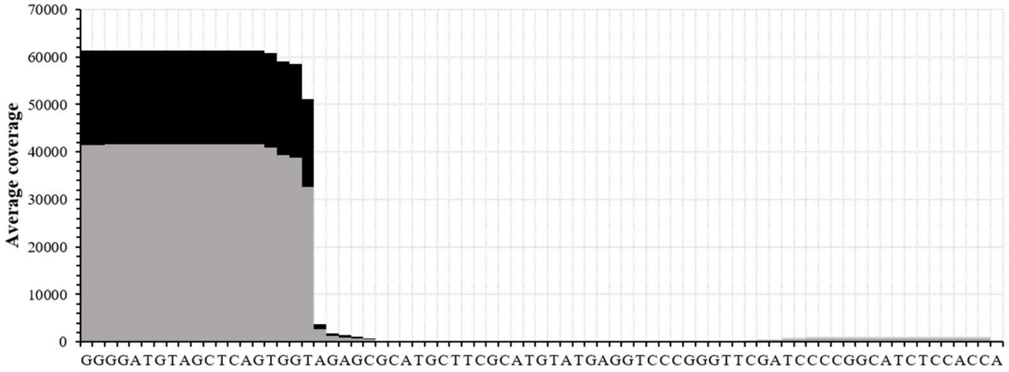 |

<table>
<tr><td>tRNA-Ala-CGC-2</td><td>0.53</td><td>0.0239</td><td>GU AG</td><td>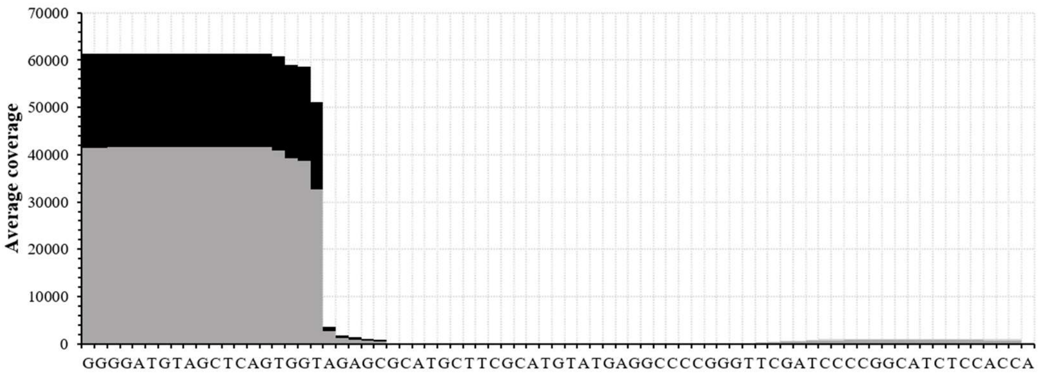
</td></tr>
<tr><td>tRNA-Ala-CGC-3</td><td>0.53</td><td>0.0240</td><td>GU AG</td><td>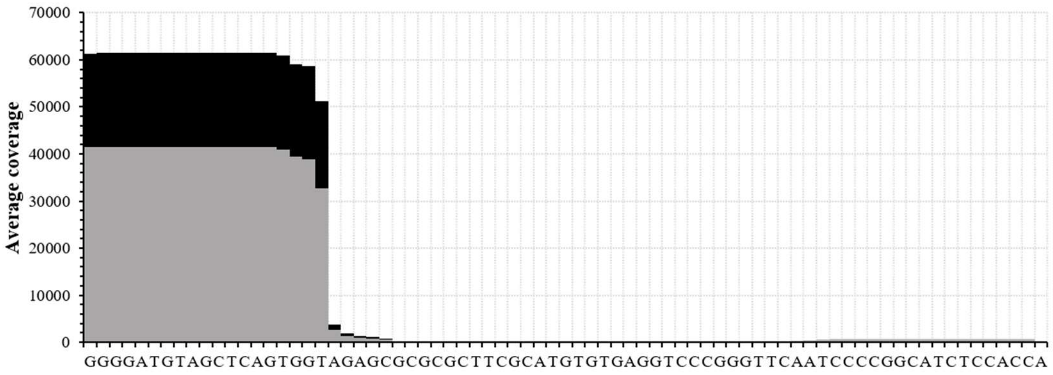
</td></tr>
<tr><td>tRNA-Ala-TGC-2</td><td>0.53</td><td>0.0239</td><td>GU AG</td><td>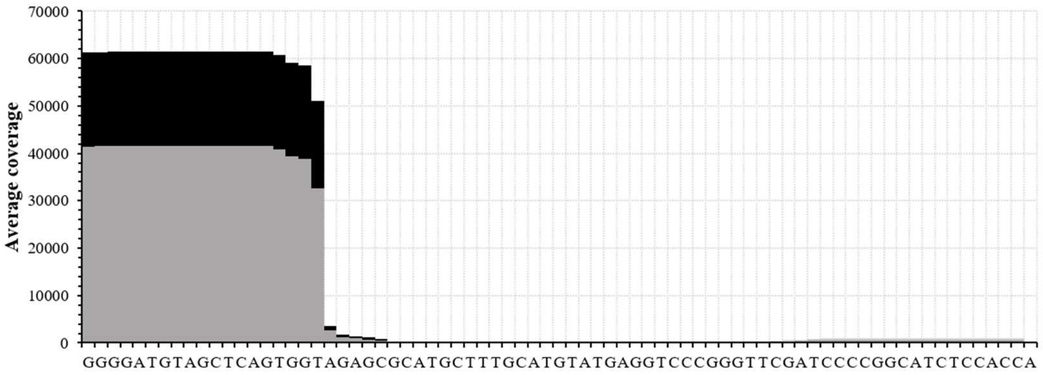
</td></tr>
</table>

<table>
<tr><td>tRNA-Ala-TGC-3</td><td>0.53</td><td>0.0239</td><td>GU AG</td><td>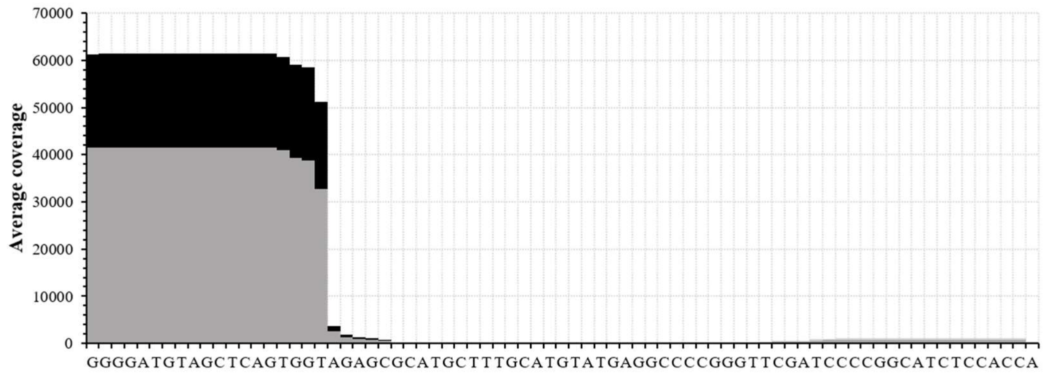
</td></tr>
<tr><td>tRNA-Ala-TGC-4</td><td>0.53</td><td>0.0235</td><td>GU AG</td><td>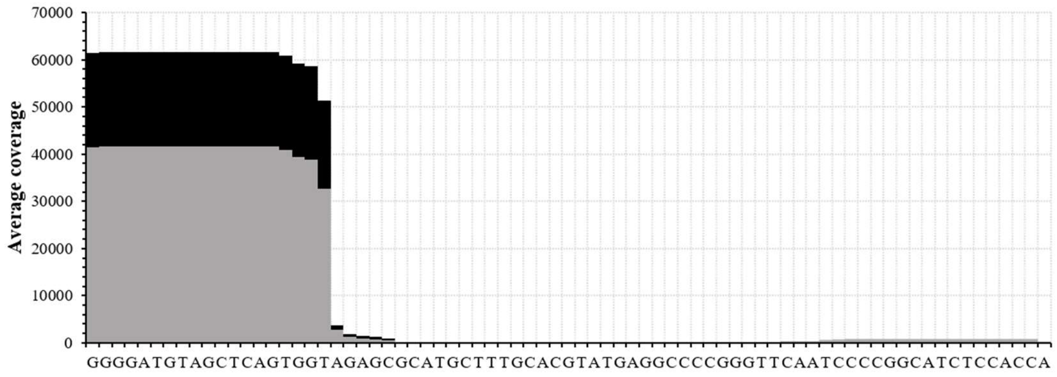
</td></tr>
<tr><td rowspan="2">tRNA-Asn-GTT-8</td><td>0.60</td><td>0.0159</td><td>GA UC</td><td rowspan="2">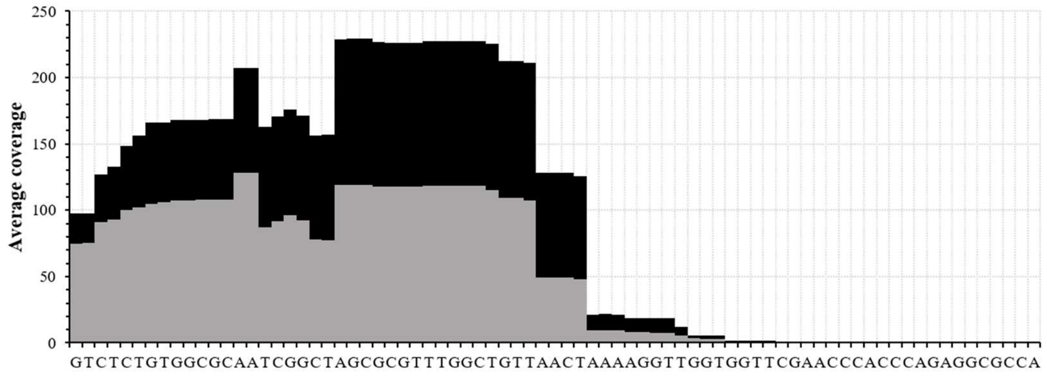
</td></tr>
<tr><td>0.73</td><td>0.0105</td><td>CU AA</td></tr>
</table>

<table>
<tr><td rowspan="2">tRNA-Asn-GTT-10</td><td>0.62</td><td>0.0123</td><td>GA|UC</td><td rowspan="2">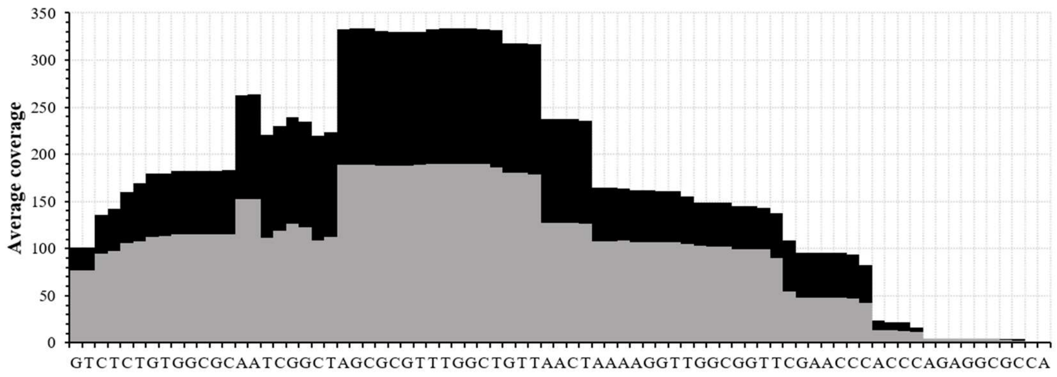
</td></tr>
<tr><td>0.62</td><td>0.0237</td><td>CU|AA</td></tr>
<tr><td rowspan="2">tRNA-Asn-GTT-12</td><td>0.60</td><td>0.0157</td><td>GA|UC</td><td rowspan="2">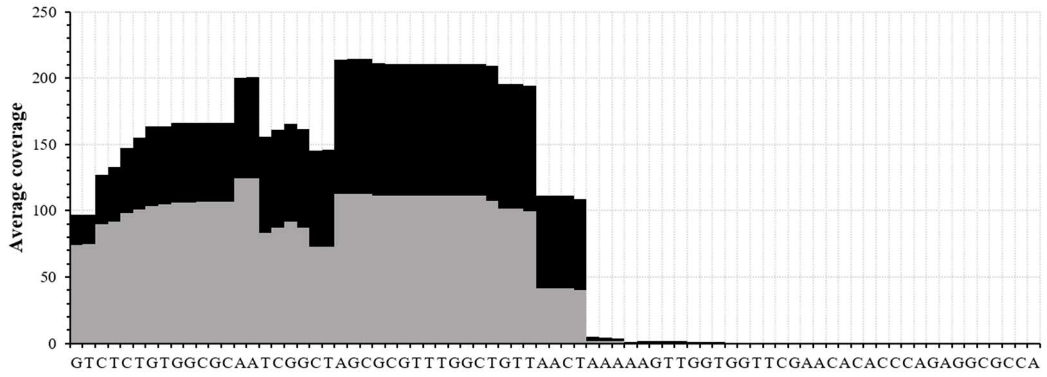
</td></tr>
<tr><td>0.70</td><td>0.0153</td><td>CU|AA</td></tr>
<tr><td>tRNA-Asp-GTC-4</td><td>0.67</td><td>0.0286</td><td>CA|CU</td><td>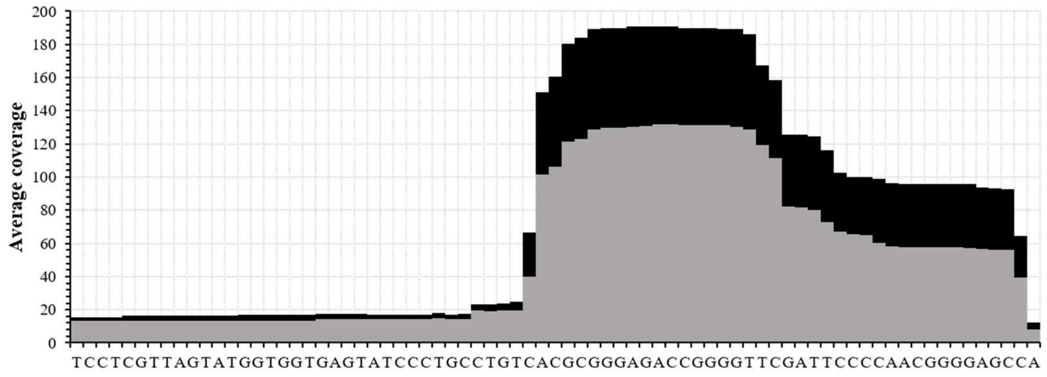
</td></tr>
</table>

| | | | | |
|---|---|---|---|---|
| tRNA-Asp-GTC-5 | 0.92 | 0.0195 | GU CA | 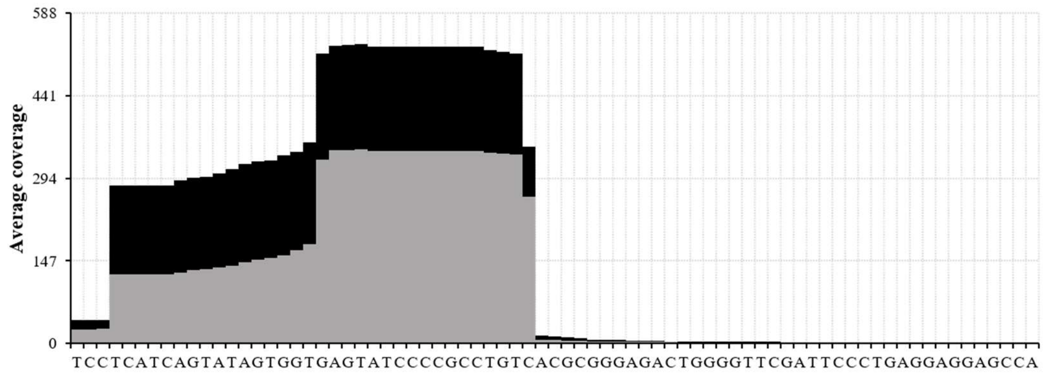  |
| tRNA-Cys-GCA-1 | 0.59 | 0.0101 | GC AG | 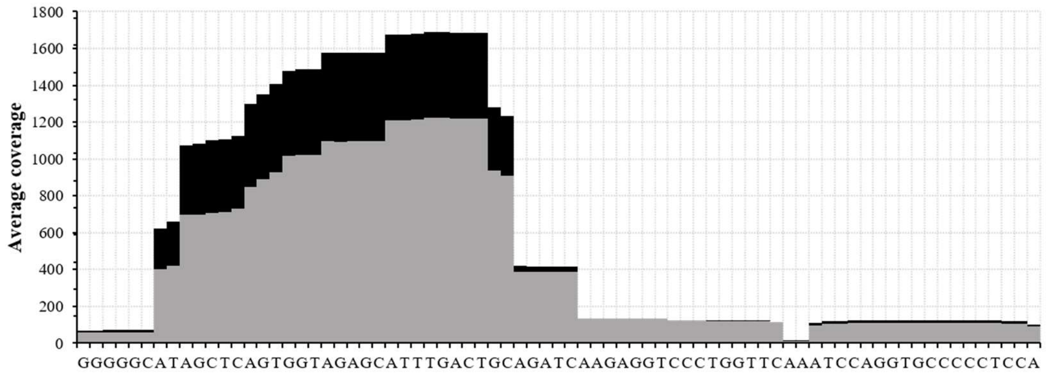  |
| tRNA-Cys-GCA-2 | 0.56 | 0.0079 | GC AG | 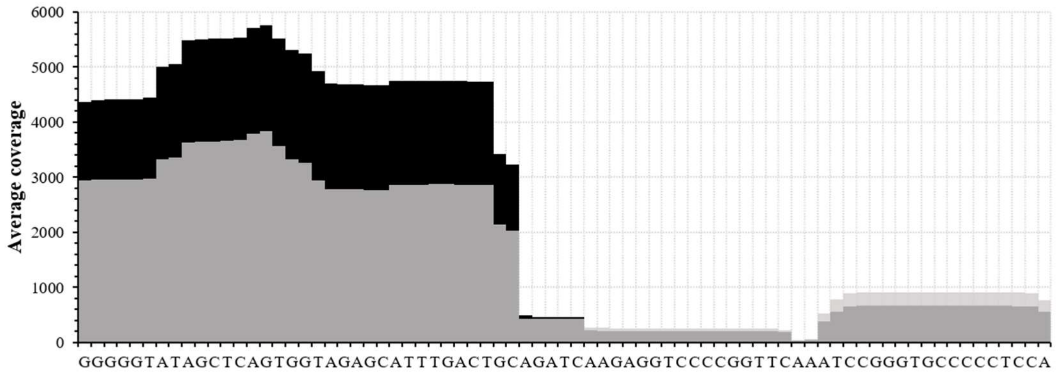  |

| | | | | |
|---|---|---|---|---|
| tRNA-Cys-GCA-4 | 0.56 | 0.0077 | GC\|AG | 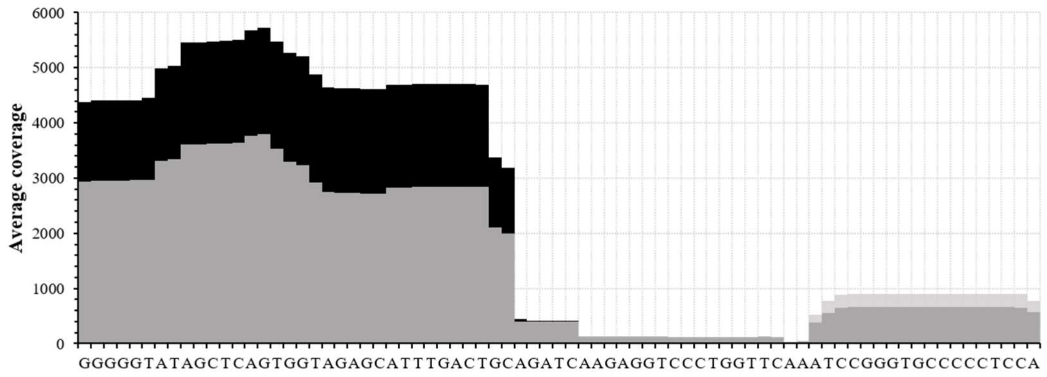 |
| tRNA-Gln-CTG-3 | 1.26 | 0.0013 | CC\|AG | 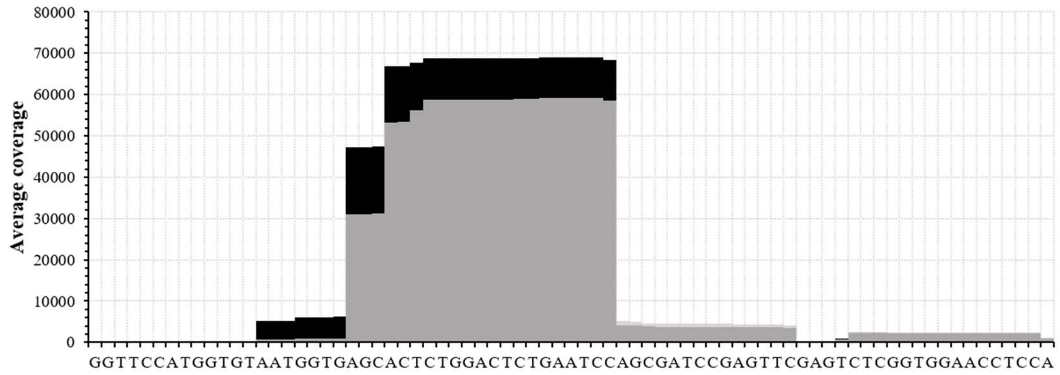 |
| tRNA-Gln-CTG-8 | 1.03 | 0.0043 | AC\|AG | 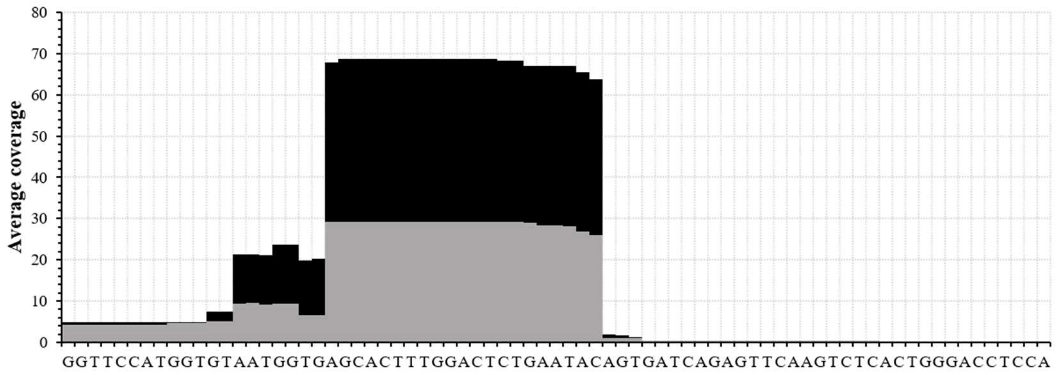 |

| | | | | |
|---|---|---|---|---|
| tRNA-Gln-TTG-1 | 1.65 | 2.6300E-08 | CC↓AG | 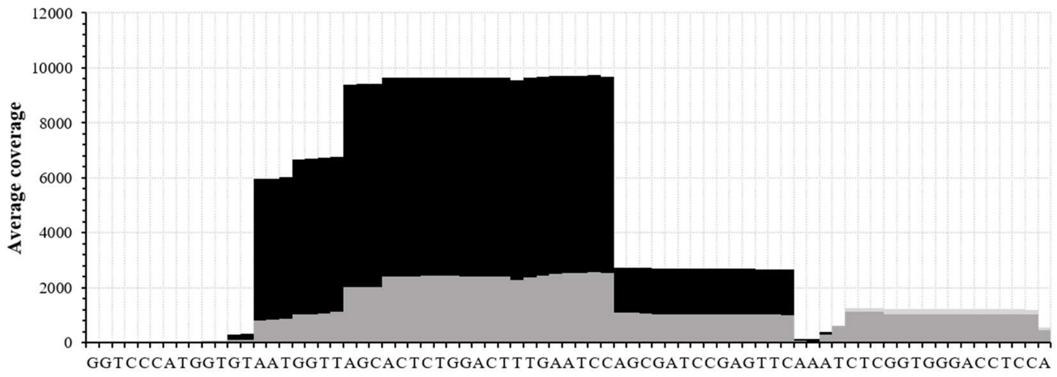 |
| tRNA-Gln-TTG-2 | 1.72 | 1.8300E-06 | CC↓AG | 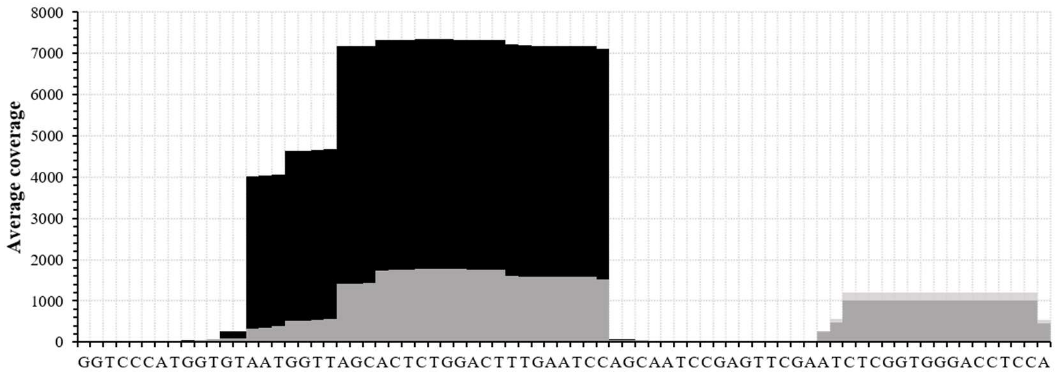 |
| tRNA-Gln-TTG-3 | 1.65 | 2.6300E-08 | CC↓AG | 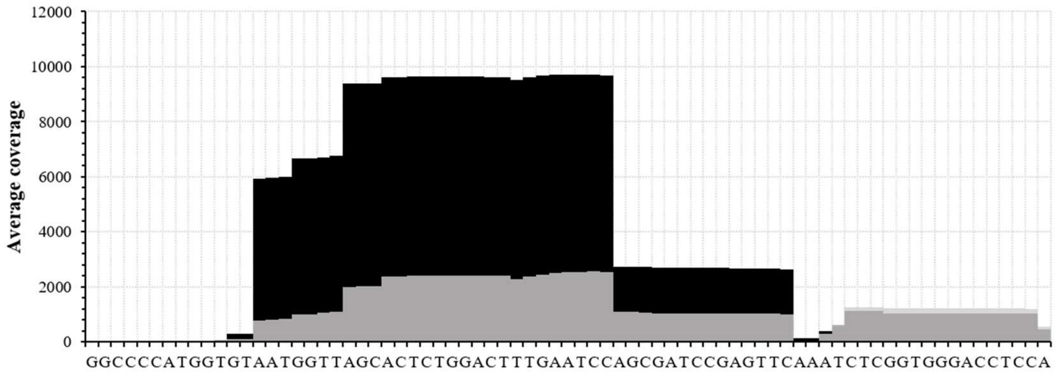 |

| | | | | |
|---|---|---|---|---|
| tRNA-Glu-TTC-4 | 0.53 | 0.0447 | CA CU | 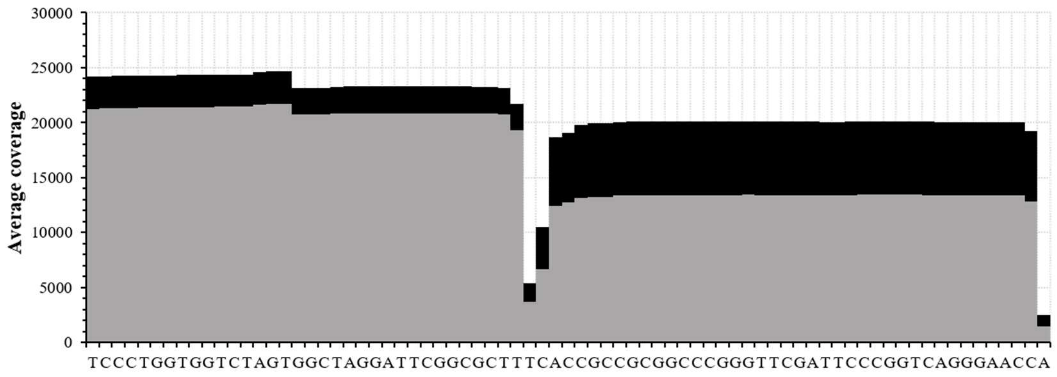 |
| tRNA-Tyr-GTA-11 | 0.63 | 0.0229 | GU AG | 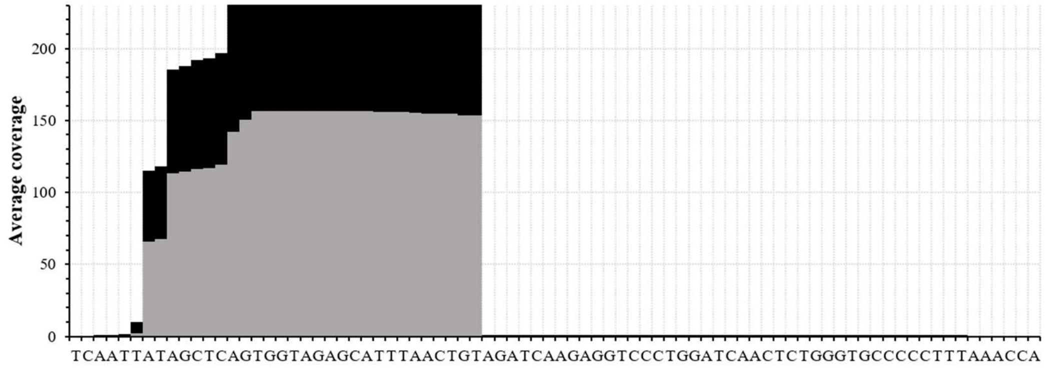 |

**S3 Table. Over-represented miRNA sequences identified in WT vs RNase2-KO THP1 cells. Putative location of cleavage sites is indicated. Information taken from miRBase.**

| Accession number | ID | $Log_2$ fold change | *P* (adjusted) | Identified Sequence | Main predicted cleavage site | Differential coverage between WT (black) and KO (grey) |
|---|---|---|---|---|---|---|
| MIMAT0000087 | hsa-miR-30a-5p | 3.04 | 2.6200E-27 | UGUAAACAUCCUCGACUGGAAG | GC↓UG | 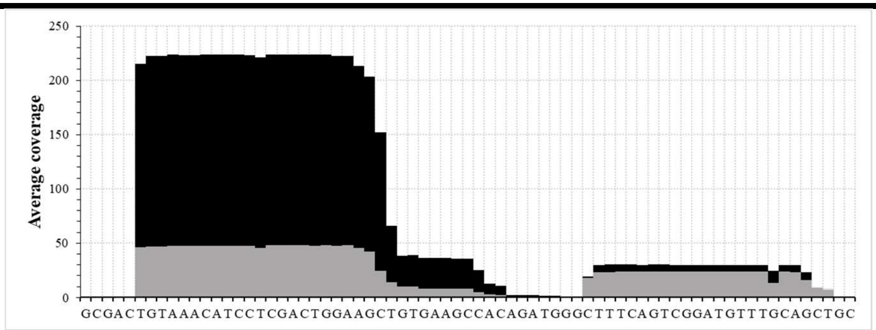 |
| MIMAT0000689 | hsa-miR-99b-5p | 2.85 | 1.5033E-21 | CACCCGUAGAACCGACCUUGCG | GC↓GG | 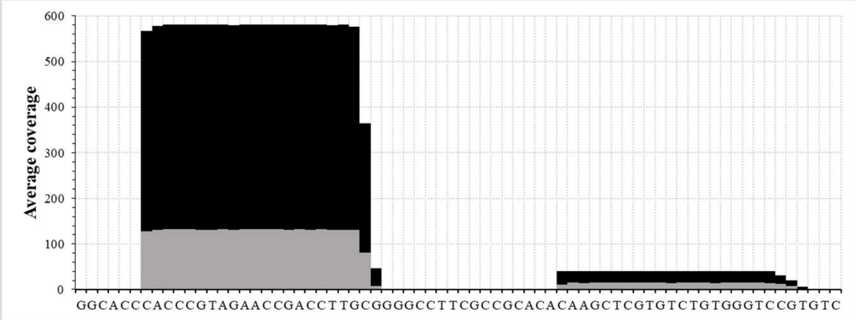 |
| MIMAT0004678 | hsa-miR-99b-3p | 2.21 | 2.7531E-04 | CAAGCUCGUGUCUGUGGGUCCG | CG↓UG | |

| | | | | | | |
|---|---|---|---|---|---|---|
| MIMAT0000098 | hsa-miR-100-5p | 2.32 | 1.4260E-19 | AACCCGUAGAUCCGAACUUGUG | GU↓GG | 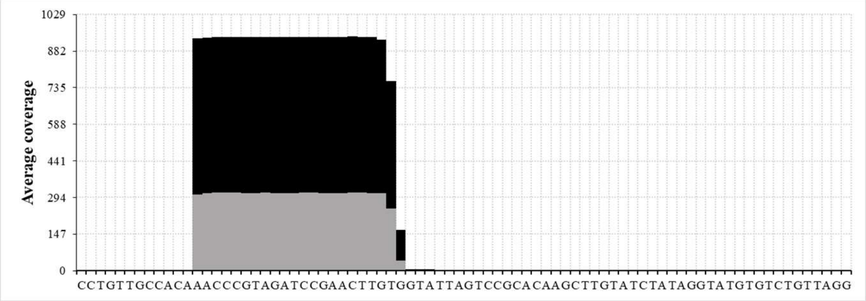 |
| MIMAT0000443 | hsa-miR-125a-5p | 3.33 | 7.8595E-15 | UCCCUGAGACCCUUUAACCUGUGA | UG↓AG | 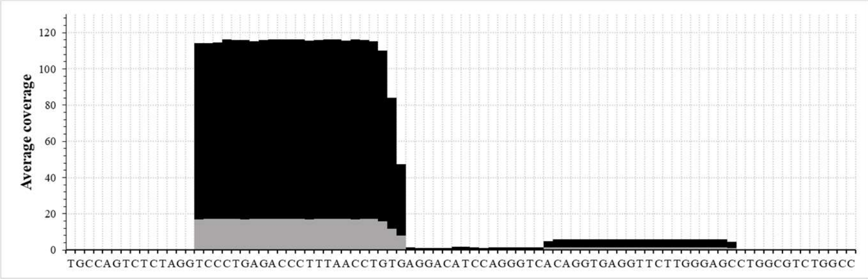 |
| MIMAT0000066 | hsa-let-7e-5p | 2.48 | 4.9694E-10 | UGAGGUAGGAGGUUGUAUAGUU | GU↓UG | 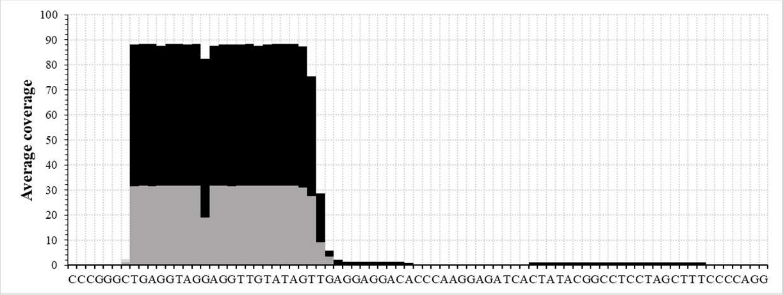 |

<table>
<tr><td rowspan="2">MIMAT00<br>00423</td><td rowspan="2">hsa-miR-<br>125b-5p</td><td rowspan="2">2.40</td><td rowspan="2">6.0555E-05</td><td rowspan="2">UCCCUG<br>AGACCC<br>UAACUU<br>GUGA</td><td>UG AU</td><td>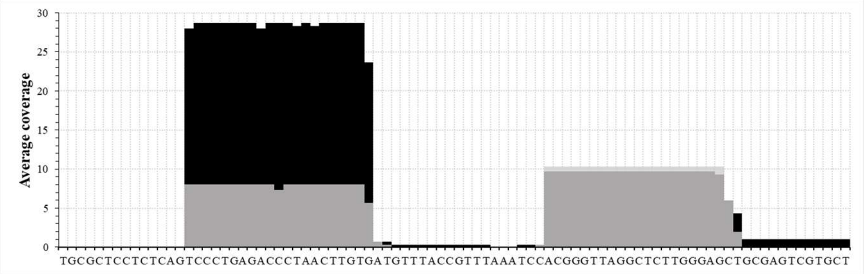
</td></tr>
<tr><td>UG AG</td><td>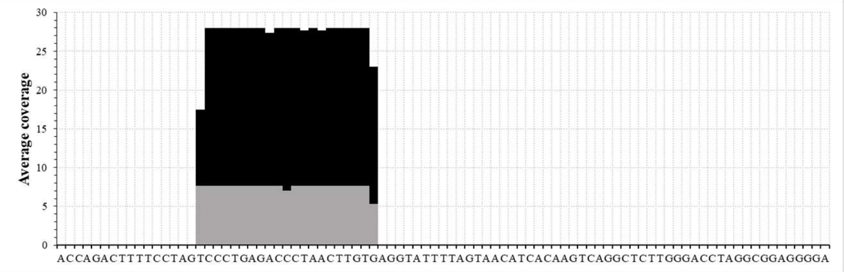
</td></tr>
<tr><td>MIMAT00<br>03150</td><td>hsa-miR-<br>455-5p</td><td>2.76</td><td>3.8997E-03</td><td>UAUGUG<br>CCUUUG<br>GACUAC<br>AUCG</td><td>GU GG</td><td>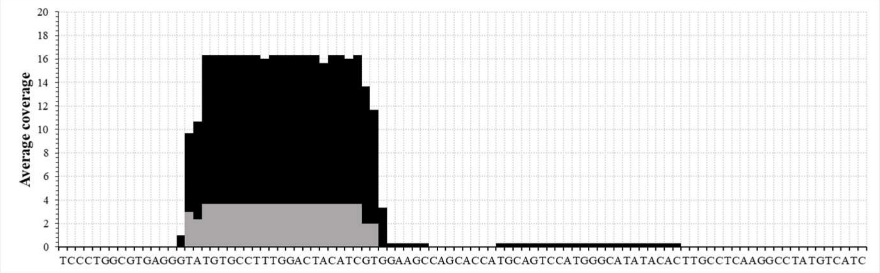
</td></tr>
</table>

**S4 Table. List of tiRNA & tRFs significantly overrepresented in WT vs RNase2-KO THP1 cells. The identified sequences and potential cleavage regions are shown.**

| Condition | Transcript name | Parental tRNA | Type | Identified sequence in tiRNA&tRF library | Remaining sequence from parental tRNA sequence | potential cleavage region[1] | tRNA cleavage region location |
|---|---|---|---|---|---|---|---|
| WT vs KO | 3'tiR_088_LysCTT (n) | LysCTT | 3-half | CUUAAUCUCAGGGUCGUGGGUUCGAGCCCCACGUUGGGCGCCA | GCCCGGCUAGCUCAGUCGGUAGAGCAUGGGACU | GGA(CUCUUAA) | Anticodon loop |
| | tiRNA-5033-LysTTT-1 | LysTTT | 5-half | GCCCGGAUAGCUCAGUCGGUAGAGCAUCAGACU | UUUAAUCUGAGGGUCCAGGGUUCAAGUCCCUGUUCGGGCG | AGA(CUUUUAA) | Anticodon loop |
| | 1039 | Pre-ArgCCT | tRF-1 | UCGAGAGGGGCUGUGCUCGCAAGGUUUCUUU | CCAAAGCA | AAGCAUCGAG | Acceptor stem |
| | 3002A | ProCGG, AGG, TGG, CGG | tRF-3 | AUCCCGGACGAGCCCCCA | GGCUCGUUGGUCUAGGGGUGUGGUUCUCGCUUAGGGCGGGAGACCCAAGAGGUCCCGGGUUCAA | (UUCAAAU)CCC | T-Loop |
| | 5028/29A | GluTCC | tRF-5 | UCCCACAUGGUCUAGCGG | UUAGGAUUCCUGGUUUUCACCCAGGUGGCCCGGGUUCGACUCCCGGUAUGGGAA | (AGCGGUUA)GG | D-loop |

| | | | | | | | |
|---|---|---|---|---|---|---|---|
| KO+RSV vs WT+RSV | 3'tiR_060_ MetCAT (n) | MetCAT | 3-half | UCAUAAUCUGA AGGUCGUGAGU UCGAUCCUCAC ACGGGGCACCA | GCCCUCUUAGUGC AGCUGGCAGCGC GUCAGUU | CAGU(UUCAUAA) | Anticodon loop |
| | TRF62 | MetCAT | itRF | AGCAGAGUGGC GCAGCGGAAGC GUGCUGG | GCCCAUAACCCAG AGGUCGAUGGAU CGAAACCAUCCUC UGCUA | GCUGGG(CCCAUAA) | D loop |
| | TRF315 | LysCTT | itRF | GCCCGGCUAGC UCAGUCGGUAG AGCAUGG | GACUCUUAAUCU CAGGGUCGUGGG UUCGAGCCCCACG UUGGGCG | CAUGGGA(CUCUUAA) | D loop |
| | TRF353 | GlnTTG | itRF | GGCCCCAUGGU GUAAUGGUUAG CACUCUGGA | CUUUGAAUCCAG CAAUCCGAGUUC GAAUCUCGGUGG GACCU | CUGGA(CUUUGAA) | D loop |
| | TRF419 | LeuTAG | itRF | GGUAGUGUGGC CGAGCGGUCUA AGGCGCUG | GAUUUAGGCUCC AGUCUCUUCGGA GGCGUGGGUUCG AAUCCCACCGCUG CCA | CGCUGGA(UUUAGGC) | D loop |
| | tiRNA-5030-LysCTT-2 | LysCTT | 5-half | GCCCGGCUAGC UCAGUCGGUAG AGCAUGGG | ACUCUUAAUCUC AGGGUCGUGGGU UCGAGCCCCACGU UGGGCG | AUGGGA(CUCUUAA) | Stem near Anticodon loop |
| | tiRNA-5031-HisGTG-1 | HisGTG | 5-half | GCCGUGAUCGU AUAGUGGUUAG UACUCUGCG | UUGUGGCCGCAG CAACCUCGGUUCG AAUCCGAGUCAC GGCA | CUGCG(UUGUGGC) | Stem near Anticodon loop |

| | | | | | |
|---|---|---|---|---|---|
| tiRNA-5031-GluCTC-1 | GluCTC | 5-half | UCCCUGGUGGUCUAGUGGUUAGGAUUCGGCG | CUCUCACCGCCGCGGCCCGGGUUCGAUUCCCGGUCAGGAAA CUCUCACCGCCGCGGCCCGGGUUCGAUUCCCGGUCAGGGAA | CGGCG(CUCUCAC) | Stem near Anticodon loop |
| 1001 | Pre-SerTGA | tRF-1 | GAAGCGGGUGCUCUUAUUUU | CUCGCUGCG | CUGCGGAAGC | Acceptor stem |
| 1013 | Pre-AlaCGC | tRF-1 | GGCGAUCACGUAGAUUUU | UCUCCA | CUCCAGGCGA | Acceptor stem |
| 1039 | Pre-ArgCCT | tRF-1 | UCGAGAGGGGCUGUGCUCGCAAGGUUUCUUU | CTGGGGU | GGGGUUCGAG | Acceptor stem |
| 3004B | GlnTTG | tRF-3 | UCAAAUCUCGGUGGGACCUCCA | GGCCCCAUGGUGUAAUGGUUAGCACUCUGGACUUUGAAUCCAGCGAUCCGAGU | CGAGU(UCAAAU) | T loop |
| 3006B | LysTTT | tRF-3 | UCAAGUCCCUGUUCGGGCGCCA | GCCCGGAUAGCUCAGUCGGUAGAGCAUCAGACUUUUAAUCUGAGGGUCCGGGGU | GGGG(UUCAAGU) | T loop |
| 3016/18/22B | LysCTT/MetCAT/IleAAT | tRF-3 | UCGAGCCCCACGUUGGGCGCCA | GCCCGGCUAGCUCAGUCGGUAGAGCAUGGGACUCUUAAUCUCAGGGUCGUGGGU | UGGG(UUCGAGC) | T loop |

| | | | | | | |
|---|---|---|---|---|---|---|
| 5024A | LeuTAA | tRF-5 | GUUAAGAUGGC AGA | ACCUGGCAGUUU CAUAAAACUUAA AGUUUAUAAUCA GAGGUUCAACUC CUCUUCUUAACA | GCAG(AACCUGGCAG ) | D loop |
| 5032A | AspGTC | tRF-5 | UCCUCGUUAGU AUAGUGG | UGAGUGUCCCCG UCUGUCACGCGG GAGACCGGGGUU CGAUUCCCCGACG GGGAG | (AGUGGUGA)GU | D loop |
| TRF356/35 9 | ArgCCG /ArgTC G | tRF-5 | GGCCGCGUGGC CUAAUGGA | UAAGGCGUCUGA CUUCGGAUCAGA AGAUUGCAGGUU CGAGUCCUGCCGC GGUCG | (AAUGGAUA)AGG | D loop |
| TRF366 | Thr-TGT | tRF-5 | GGCUCCAUAGC UCAGUGGUUAG AGCA | CUGGUCUUGUAA ACCAGGGGUCGC GAGUUCGAUCCU CGCUGGGGCCU | (AGUGGUUA)GAGCA CUGGU | D loop |
| TRF365 | Thr-TGT | tRF-5 | GGCUCCAUAGC UCAGGGGU | UAGAGCGCUGGU CUUGUAAACCAG GGGUCGCGAGUU CAAUUCUCGCUG GGGCCU | (AGGGGUUA)GAG | D loop |
| TRF396 | AlaAGC | tRF-5 | GGGGGUAUAGC UCAGCGGU | AGAGCGCGUGCU UAGCAUGCACGA GGUCCUGGGUUC AAUCCCCAAUACC UCCA | (AGCGGUA)GAGC | D loop |

| | | | | | | | |
|---|---|---|---|---|---|---|---|
| | TRF457 | SerAGA/ SrTGA | tRF-5 | GUAGUCGUGGC CGAGUGG | UUAAGGCGAUGG ACUAGAAAUCCA UUGGGGUUUCCC CACGCAGGUUCG AAUCCUGCCGACU ACG | (GAGUGGUUAA)G | D loop |
| | TRF550/55 1 | | tRF-5 | UCCUUGGUGGU CUAGUGGCUAG | GAUUCGGCGCUU UCACCGCCGCGGC CCGGGUUCGAUU CCCGGCCAGGGAA | (AGUGGCUA)GGAUU C | D loop |
| RSV vs WT | tiRNA-5034-ValCAC-3 | ValCAC | 5-half | GUUUCCGUAGU GUAGCGGUUAU CACAUUCGCCU C | ACACGCGAAAGG UCCCCGGUUUGA AACCAGGCGGAA ACA | GC(CUCACAC)G | Anticodon loop |
| | tiRNA-5029-ProAGG | ProAGG | 5-half | GGCUCGUUGGU CUAGGGGUAUG AUUCUCG | CUUAGGGUGCGA GAGGUCCCGGGU UCAAAUCCCGGAC GAGCCC | UCUCGC(UUAGGGU) | Anticodon loop |
| | TRF354 | ThrTGT | tRF-5 | GGCCCUAUAGC UCAGGGG | UUAGAGCACUGG UCUUGUAAACCA GGGGUCGCGAGU UCAAAUCUCGCU GGGGCCU | (AGGGGUUA)GA | D loop |
| | TRF374 | ThrCGT | tRF-5 | GGCUCUGUGGC UUAGUUGGC | UAAAGCGCCUGU CUCGUAAACAGG AGAUCCUGGGUU CGAAUCCCAGCGG GGCCU | (AGUUGGCUA)AAG | D loop |

[1]the potential cleavage region of the parental tRNAs are inferred according to the sequences detected in the tiRNA&tRFs library screening. Naming is directly taken from the tiRNA&tRFs library. Complementarily, we include in the next column the remaining tRNA sequence from the parental tRNA. The first 5 bases from tRNA fragments at the 5'

and 3' sides are coloured in red and blue, respectively. The full region corresponding to specific loops in the parental tRNA is included and enclosed by parenthesis. When more than one sequence is found for the same identifier, the first one is taken (in no case, the punctual base differences are affecting the cleavage region). Information for tRNAs sequences was taken from GtRNAdb, MINTbase, tRNAdb and tRFdb databases.

**S5 Table. Comparison of cleavage specificity for RNA synthetic substrates *in vitro* and ncRNA in cell assays for other RNaseA superfamily members**.

| RNase | Synthetic substrate | Base preference | Ref. | ncRNA | Base preference | Ref. |
|---|---|---|---|---|---|---|
| **RNase5** | | **B1** | | | | |
| | CpA/UpA | C/U ~12 | 1,2 | Selected tRNA (anticodon loops) | | 3 |
| | | | | CA and CU (anticodon loops) | | 4 |
| | | **B2** | | | | |
| | CpA/CpG | A/G ~3 | 1,2 | | | |
| | CpG/CpC | G/C ~3 | 1 | | | |
| | | | | tRNA (Anticodon loops) | A>U | 4 |
| ***Rana pipiens* RNase (Onconase)** | | **B1** | | | | |
| | UpG/CpG | U/C ~10 | 5 | tRNA | Major: UG↓G (Variable loop and D arm) | 6, 7 |
| | | | | tRNA | Minor: G↓U C↓U | 6,7 |
| | polyU/polyC | U/C ~100 | 8 | | | |
| | tetranucleotide | U/C ~60 | 8 | | | |
| | | **B2** | | | | |
| | Tetranucleotide | G/A ~800 | 8 | | | |

**S6 Table. Primers used for PCR.**

| Name | Sequences |
|---|---|
| Short-Fw | CCACCGGGTCTTCGAAGACTGTTTG |
| Short-Rv | CATGGGTGGCCCAGAACCTTCTGACAAACGATC |
| Cherry-Kosak-AgeI | TATAACCGGTGCCACCATGGTGAGCAAGGGCGAGGA |
| Cherry-BamHI | ATAAGGATCCCTTGTACAGCTCGTCCATGC |
| GFP-BsrGI | ATATTGTACAGAGGGCAGAGGAAGTCTGCT |
| GFP-EcoRI | TTAGAATTCTTACAGCTCGTCCATGCCGAGAG |
| RSV-A-Fw | CTCAATTTCCTCACTTCTCCAGTGT |
| RSV-A-Rv | CTTGATTCCTCGGTGTACCTCTGT |
| RSV-A-Probe | TCCCATTATGCCTAGGCCAGCAGCA |

The fluorogenic probe was labelled with 6-carboxyfluorescein (FAM) and 6-carboxytetramethylrhodamine (TAMRA).
